# Radiogenomics-based Risk Prediction of Glioblastoma Multiforme with Clinical Relevance

**DOI:** 10.1101/350934

**Authors:** Xiaohua Qian, Hua Tan, Wei Chen, Weiling Zhao, Michael D. Chan, Xiaobo Zhou

**Author notes:** Co-first authors (Xiaohua Qian, Hua Tan). Corresponding to: Xiaobo Zhou, School of Biomedical Informatics, the University of Texas Health Science Center at Houston, Houston, TX 77030, USA.

## Abstract

GBM is the most common and aggressive primary brain tumor. Although the TMZ-based radiochemotherapy improves overall GBM patients’ survival, it also increases the frequency of false positive post-treatment magnetic resonance imaging (MRI) assessments for tumor progression. Pseudoprogression is a treatment-related reaction with an increase in contrast-enhancing lesion size at the tumor site or resection margins which mimics tumor recurrence on MRI. Accurate and reliable prognostication of GBM progression is urgently needed in the clinical management of GBM patients. Clinical data analysis indicates that the patients with PsP had superior overall and progression-free survival rates. In this study, we aimed to develop a prognostic model to evaluate tumor progression potential of GBM patients following standard therapies. We applied a dictionary learning scheme to obtain imaging features of GBM patients with PsP or TTP from the Wake dataset. Based on these radiographic features, we then conducted radiogenomics analysis to identify the significantly associated genes. These significantly associated genes were then used as features to construct a 2YS logistic regression model. GBM patients were classified into low-and high-survival risk groups based on the individual 2YS scores derived from this model. We tested our model using an independent TCGA dataset and found that 2YS scores were significantly associated with the patients’ overall survival. We further used two cohorts of the TCGA data to train and test our model. Our results show that 2YS scores-based classification results from the training and testing TGCA datasets were significantly associated with the overall survival of patients. We also analyzed the survival prediction ability of other clinical factors (gender, age, KPS, normal cell ratio) and found that these factors were not related or weakly correlated with patients’ survival. Overall, our studies have demonstrated the effectiveness and robustness of the 2YS model in predicting clinical outcomes of GBM patients after standard therapies.

## Introduction

Glioblastoma multiforme (GBM) is the most common and aggressive primary brain tumor in adults, with the median survival of 14–16 months and an average 2-year survival rate of 26–33%^1^. The current standard of care is surgical resection followed by radiotherapy and adjuvant chemotherapy with temozolomide (TMZ)^2^. Although radiochemotherapy with TMZ is superior to radiotherapy alone in the treatment of GBM, this intensified treatment leads to an increased rate in pseudoprogression (PsP)^3^. PsP is a subacute and post-treatment reaction of imaging changes at the tumor site or resection margins. The contrast enhancement and considerable vasogenic edema on the conventional postoperative monitoring MRI can mimic early tumor progression, making early diagnosis difficult for the GBM patient through the current imaging techniques, although these imaging changes subsequently regress or remain stable^4,5^.

Although pathological confirmation is the reference standard in clinical management of GBM treament; however, it is not desirable in clinical practice due to the requirement of a second surgery. Follow-up imaging has been employed to make a diagnosis based on the morphological changes of suspected lesions. To achieve a satisfactory diagnostic accuracy, it usually takes several months in the current clinical practice, thus affecting the clinical management of GBM patients. Therefore, early diagnosis and prediction of PsP are critically important for the prognosis of GBM patients. Many efforts have also been devoted to exploring imaging signatures or genomics biomarkers for the diagnosis or prediction of clinical outcome of GBM patients. Perfusion parameters, especially tumor blood volume, have been used as a prognostic signature for disease progression or survival^6–9^. Other morphologic imaging features from the non-enhanced and contrast-enhanced tumors, necrosis, and edema also serve as prognostic markers of GBM^10,11^. Several genomics biomarkers, such as MGMT promoter methylation^12^, Ki67 expression^13^, IDH1 mutation^14^ and p53 mutation^15^, have been associated with the development of PsP in GBM patients. Among these biomarkers, MGMT promoter methylation has attracted a lot of attention^3,12,13,16–18^. However, its predictive potential in PsP remains debatable^14,19–22^. Currently, there are no systematical and comprehensive approaches for accurate stratification of PsP and TTP and prediction of clinical outcomes of GBM patients receiving standard treatment. According to clinical data analysis, the patients with PsP had superior overall and progression-free survival rates^23,24^. However, most of previous studies did not associate imaging signatures or genetic biomarkers with patient survival. Interestingly, we analyzed the private dataset in our institute and found that there was no significant difference in patients’ survival between PsP and TTP groups diagnosed by clinicians, highlighting the difficulty of obtaining accurate diagnosis using conventional clinical methods. An incorrect diagnosis of a PsP could result in erroneous termination of an effective treatment with a potentially negative influence on patients’ survival. Therefore, a novel and reliable diagnostic and predictive risk model with clinical relevance is required for distinguishing these two groups of GBM patients.

Radiogenomics research mainly refers to the relationship between the patient genetics and imaging characteristics^25^. Exploring this relationship will be useful for accurately assessing the risk of GBM patients in the development of PsP or TTP. However, comprehensive analysis imaging signatures and genomics biomarkers to construct a predictive survival risk model is still challenging. For example, Beck et al. found the prognostic model score from the image-based model was strongly associated with patients’ overall survival, but independent of clinical, pathological, and molecular factors of breast cancer^26^. Additionally, gene expression profiles may provide more predictive power of the disease outcome than standard systems based on clinical and histologic criteria^27,28^.

In this study, we developed a novel radiogenomics-based 2-year survival (2YS) risk predictive model using machine learning approach to stratify patients into high and low survival risk groups. The radiogenomics study was used to integrate the imaging characteristics and genetic profiles of patients for determining the significant genes and 2YS model for stratify GBM patients into survival-associated groups. For the radiogenomics study, we applied dictionary learning method and feature selection scheme to derive the discriminative imaging features from diffusion tensor imaging (DTI) of PsP and TTP. We then explored the relationship between the identified imaging features and differentially expressed genes of these two groups using the sparse regression model. The genes most relevant to the imaging features were considered as the significant genes. We then applied the significant genes as features to build a 2YS predictive model. The patients were divided into high and low-survival risk groups based on their 2YS scores. Our predictive model was developed based on the dataset from Wake Forest University and independently validated using the public dataset of TCGA. The 2YS as a measure is independent of clinical indicators, such as age, race, Karnofsky Performance Scale (KPS), but is associated with the IDH1 mutation. The major advantage of this work is the combination of imaging characteristics and gene expression data with patients’ survival to discriminate PsP and TTP. Therefore, our developed 2YS predictive model is expected to be a powerful tool for personalized diagnosis and care for GBM patients.

## Materials and Method

### The pipeline of 2YS model

The Fig. 1 shows the overview of the radiogenomics study pipeline and building procedure of prognostic model. The discriminative imaging features were derived from the DTI of PsP and TTP patients’ groups (Fig. 1a). We then explored the relationship between the identified imaging features and differentially expressed genes of these two treatment phenotypes using the sparse regression model. The genes most relevant to the imaging features were considered as the significant genes (Fig. 1b). The prognostic model was constructed based on the significant genes from the patients who have survived for 2 years and died in 2 year after therapy. The 2YS score was applied to stratify GBM patients into high-or low-risk groups.

**Figure 1.**
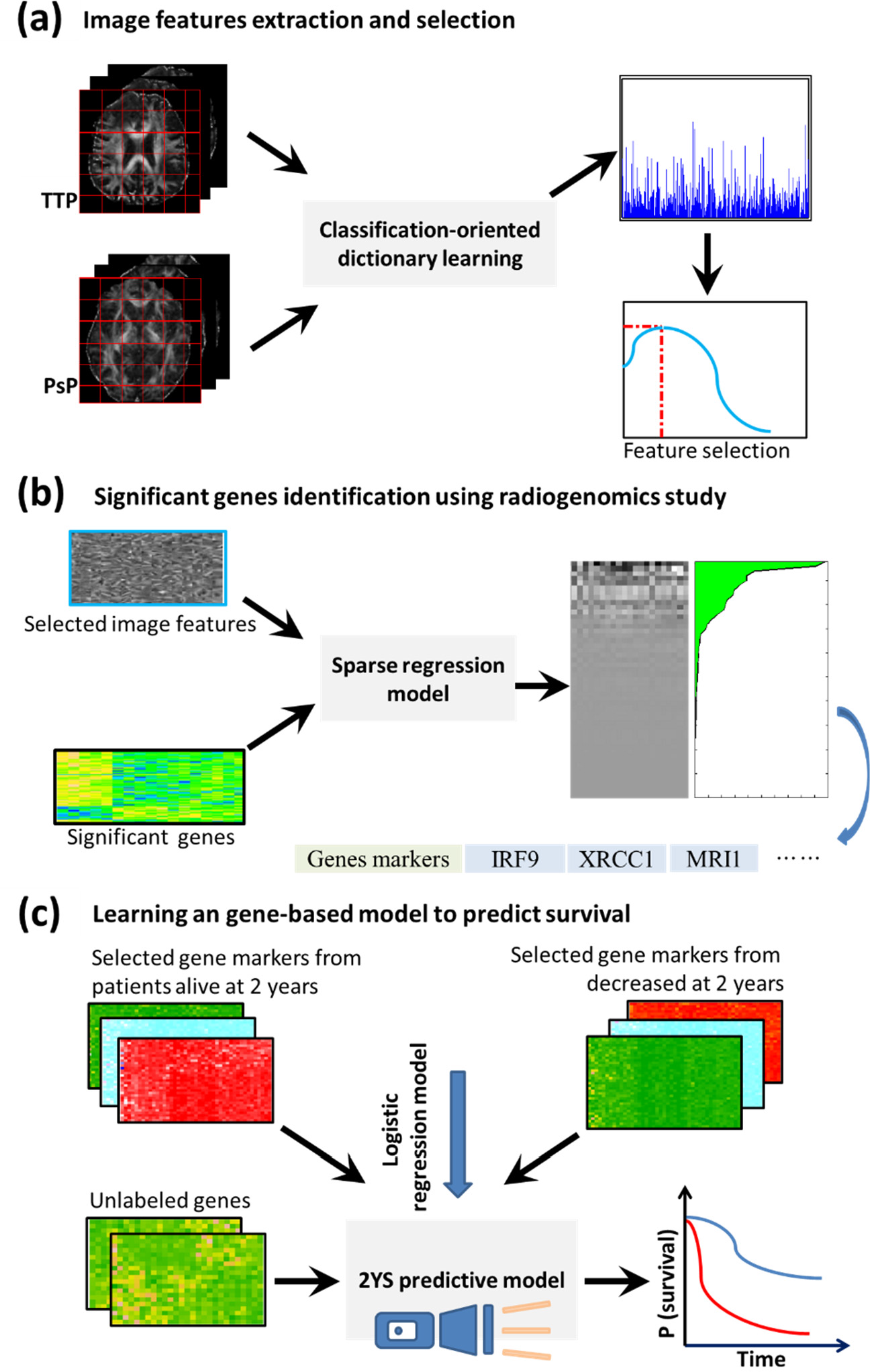
Overview of the radiogenomics study pipeline and prognostic model building procedure. This retrospective study was approved by the Institutional Review Board of Wake Forest School of Medicine. We acquired data from two independent cohorts: Wake Forest School of Medicine, namely Wake dataset, and the Cancer Genome Atlas (TCGA, http://cancergenome.nih.gov/).

### Data collection

In the Wake Forest database, more than 200 GBM patients underwent standard multimodal treatment, including surgical resection and then concurrent radiotherapy and chemotherapy with the alkylating drug TMZ. Along with radiochemotherapy, the patients underwent DTI scans (scanner: SIMCGEMR, GE Medical systems) every three months for post-surgical monitoring. The clinical diagnosis for PsP and TTP was mainly dependent on morphological changes of lesions on the follow-up imaging and clinical knowledge of the physicians. We retrospectively selected DTI scans for the patients diagnosed with PsP or TTP. As a result, we collected both DTI data and electronic clinical records for 84 patients (23 with PsP and 61 with TTP). Genetic data were collected from 42 patients and profiled using Affymetrix HuEx-1_0-st-v2 array, Illumina HumanMethylation450 BeadChip, and Affymetrix whole genome SNP 6.0 array. The corresponding electronic clinical records, including age, sex, date of surgery, and date of death, were reviewed to determine patient characteristics and treatment outcomes. The Wake dataset was applied to determine the significant genes as features and train the predictive model for survival.

The TCGA dataset from multi-institutions was applied to validate the survival predictor. Since gene expression data and survival information such as date of surgery and date of death were required for each patient, we totally collected 158 cases with genetic data and electronic clinical records from TCGA. Image feature extraction and selection were conducted using dictionary learning approach.

The purpose of deriving imaging features is to conduct the radiogenomics-study for determination of significant genes, thereby emphasizing their relationship with the progression of PsP or TTP. To achieve this goal, we extracted and selected discriminative features from DTI for classification of these two phenotypes, since DTI showed power in distinguishing these two phenotypes in previous studies^29,30^. However, segmentation of the suspected lesion area on DTI is a challenging task; thus, we proposed a dictionary learning strategy to capture the fine difference between two classes, avoiding the DTI segmentation. The feature extraction and selection procedures were comprised of three steps, i.e., preprocessing DTI data, discriminative dictionary learning, and feature extraction and selection.

### Preprocessing DTI data

We corrected current-induced distortions and subject movements of DTI and conducted the skull stripping to obtain the brain images. Fractional anisotropy (FA) value of each voxel in DTI was then calculated using FSL software and linearly registered to the standard brain template FMRIB58 in FSL. Identical resolution and number of slices for each registered volumetric FA data were confirmed.

### Discriminative dictionary learning

Although the FA values are lower for the lesion sites of PsP than that with TTP^29^, the difference between them is subtle. We previously developed a discriminative dictionary learning scheme^31^ to capture the subtle distinctions between PsP and TTP. Specifically, we directly learned specific dictionaries for PsP and TTP to gain classification-oriented characterizations and simultaneously learned shared dictionaries to describe the common features. Intuitively, PsP-specialized dictionaries with the shared dictionaries can outperform TTP-specialized dictionaries with shared dictionaries in the representation of the PsP data; thus, we applied a dual-sparse encoding strategy on each identical case to obtain the discriminative sparse coefficients^31^. Then, we used the sparse coefficient corresponding to the specific dictionaries to construct a new discriminative sparse matrix, which contains more discriminative information attributed to discarding the shared patterns.

### Feature extraction and selection

Sparse coefficients are the weights of different atoms in the dictionary for the representation of the data. We applied the max-pooling technique to extract the biggest sparse coefficients (i.e., biggest contributions) of each corresponding atoms in an indivial patient as imaging features from the constructed discriminative sparse matrix^31^.

Since the distinctions between PsP and TTP on DTI are subtle, we applied a DX score-based feature scoring system to determine the most relevant features^32,33^ with a high discrimination power between PsP and TTP. The effectiveness and efficiency of this feature selection method have been confirmed in our previous studies^31–33^. Briefly, *DX*_*score*_ is used to measures the degree of dissimilarity between positive and negative samples for each individual feature, normalized by the sum of variances in respective sample types, It can be mathematically defined as:

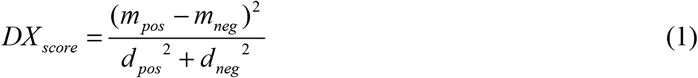

where *m*_*pos*_ and *m*_*neg*_ are the mean values of the positive and negative samples, respectively, while *d*_*pos*_ and *d*_*neg*_ are the corresponding standard deviations.

Then, we constructed a feature set by consecutively adding each feature with *DX*_*score*_ from high to low, and assessed its classification performance by 10-fold cross validation (CV) using the well-established SVM tool LIBSVM^34^. We also conducted a grid search on the radial basis function parameters (i.e., *γ* and the trade-off coefficient *C*) for each fold experiment. We identified the best accuracy and its corresponding feature set, which served as the optimal feature set.

### Identification of significant genes using radiogenomics study

To identify a compact set of significant genes closely related to the clinical treatment outcomes, we utilized a Wilcoxon rank sum test to compare the gene expression levels in the PsP and TTP groups and selected the significantly differentiated expressed genes with p-values less than 0.005. Second, we applied a sparse regression approach to reveal the associations between gene expression profiles and imaging features extracted by the dictionary learning.. Thus, we also developed a low-rank sparse regression model to effectively reduce the redundant information since different imaging features are interrelated to each other and their effects during the association process could be overlapped ^35^. According to the associated map (i.e., weight coefficients), we identified a compact set of genes whose expression values were closely related to the imaging features. Specifically, we calculated the overall weights for the expression of each gene with respect to all imaging features. The genes with top-ranked weight were selected as the significant genes. These genes were highly correlated to the imaging features.

### Construction of 2-year survival model with machine learning

We constructed a predictive model based on gene expression data to predict the 2YS score and then to divide the patients into two categories of high-and low-risk associated with survival time. We utilized the logistic regression model to build this prediction model with the input variables of significant genes’ expression values and the output of 2YS probability of each patient. Since the complete survival records were available in this study for GBM patients, it was easy to obtain the individual 2YS probability. Consequently, this trained model was used to predict the 2YS score for each patient with gene expression data. To differentiate these GBM patients into different risk subtypes, we used exhaustive search scheme to identify a threshold for the 2YS score. Patients were classified into subtypes based on the candidate threshold and the differences between subtypes assessed by the p-value in Log-rank test. The smallest p-value (i.e., most significant survival difference between two groups) in all of these subtypes was determined, and the corresponding candidate threshold was chosen as the optimal threshold for this model. By this way, we built the 2YS predictive model and determined the threshold, thereby stratifying the GBM patients into high-and low-survival risk groups. 39 cases from Wake dataset were used to train this predictive model and then we applied this model to the TCGA dataset of 158 cases from multi-institutions for validation. We also conduct the random experiments on TCGA dataset to confirm the performance of this model. Briefly, the 158 cases of TCGA dataset were randomly split into two-folds as the training and validation set. The Log-rank p-value was calculated on the classified high-and low-risk groups. Kaplan-Meier curves were applied to illustrate the survival difference.

### Analysis of the relevance between 2YS scores and clinical factors

We conducted a correlation analysis of 2YS scores and clinical factors for the Wake and TCGA datasets. GBM patients were divided into PsP or TTP based on their 2YS scores and then the correlation of individuals with each clinical variable was calculated. The trends of continuous variables (such as ages, tumor/normal cell ratio) was evaluated using Kolmogorov-Simonov test; and the trends of a discrete variable, including gender, race, MGMT, were tested by Fisher exact test. Although most of these clinical factors or measurements were identical in the Wake and the TCGA datasets, the consistency of correlations between two datasets was not emphasized here due to the diversity of clinical records in different cohorts. To assess the prognostic value of 2YS scores in the context of other clinical factors, we also conducted a multivariate Cox proportional hazards analysis for overall survival on two cohorts. Statistical significance of each variable in multivariate Cox proportional model was evaluated by calculating each variable’s Wald statistic and associated *p* value.

### Identification of significant gene features in the 2YS model

In order to identify the powerful features (i.e., genes) of the 2YS model, we conducted a bootstrap analysis of the TCGA dataset. One thousand bootstrap iterations were performed to generate 95% confidence intervals (CIs) for predicting the coefficient estimates for each gene’s expression value. If 95% CIs for an individual gene are all positive or negative, the corresponding features of this gene were considered as the significant features for 2YS predictor. Thus, the identified significant features were used to predict clinical treatment progression.

## Result

### Extraction of image features from DTI

**Figure 2.**
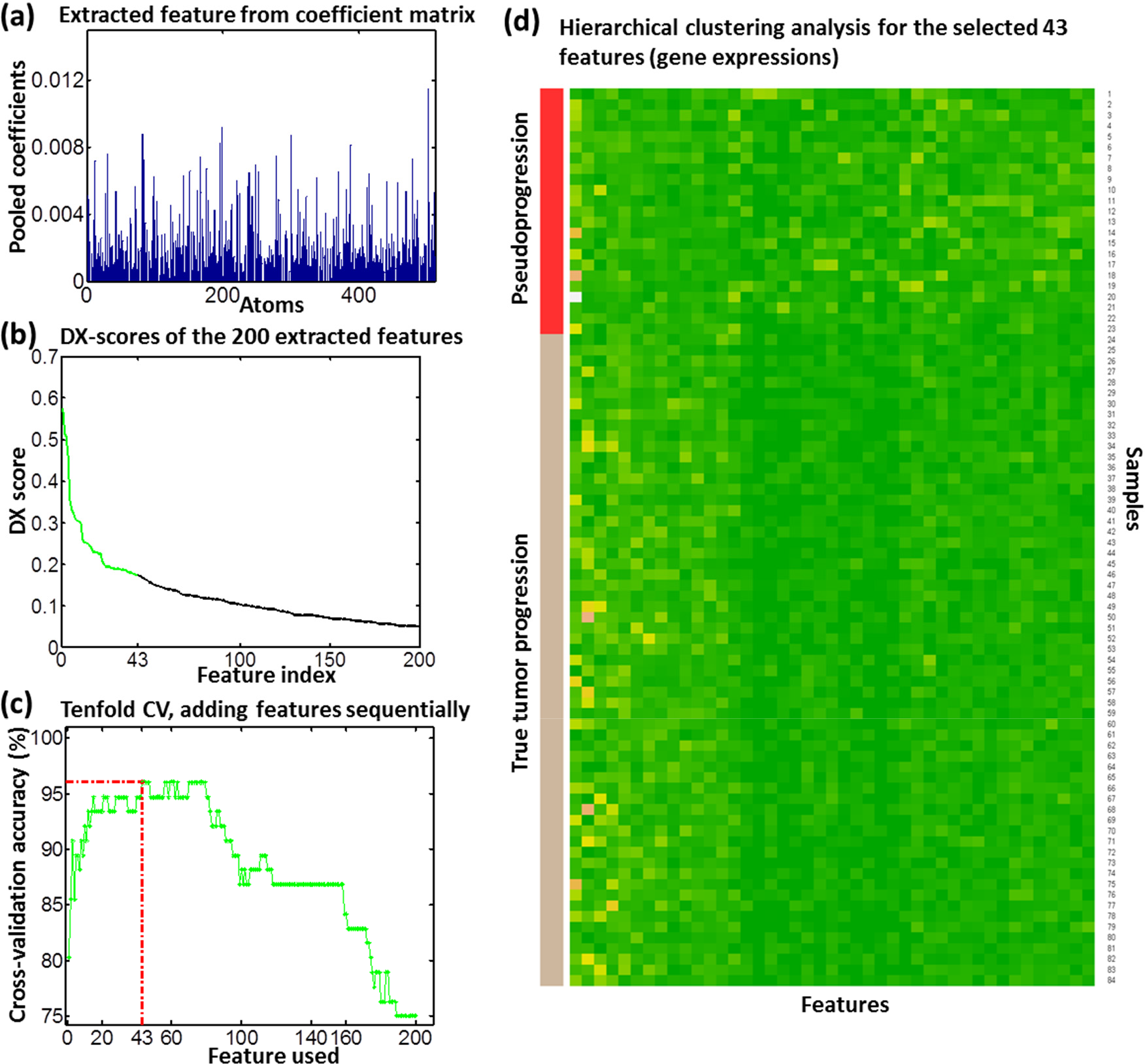
The performance of the extracted imaging features by dictionary learning of DTI. Panel a shows the extracted imaging features using max-pooling technique and b shows the DX scores of these features. Panel c shows tenfold CV analysis for the features added in one-by-one according to (b). Inclusion of the 43 top-ranked features yielded the highest CV accuracy. Panel d shows the hierarchical clustering analysis of the 43 selected features. Samples with labels from 1 to 23 are PsP cases; 24-84 are TTP cases.

To improve computational efficiency, we cropped the original resolution 256×256 of images to 164×143. We also set the following empirical parameters according to our previous studies^31^. The patch size was set as 13, PsP-and TTP-specialized dictionary sizes as 100, shared dictionary size as 10, and the number of non-zero coefficients as 10. Using the classification-oriented dictionary learning algorithm, we derived PsP- and TTP-specific dictionaries and shared dictionaries. With the dual sparse encoding scheme, we then excluded coefficients of the shared dictionaries, and used the remaining data (i.e., coefficients from the specific dictionaries) to construct a new sparse coefficient matrix. We then applied the max pooling scheme to extract features from the coefficient matrix, as shown in **Fig. 2a**. The pooled features (i.e., a histogram for all atoms) represent the greatest contributions of each corresponding atom in a specific case.

To obtain the most discriminative features, we assessed these pooled features using proposed feature scoring systems^32,33^. The pooled features were sorted by *DX*_*score*_ (**Fig.2b**), which indicates the features with higher scores have the better ability in discriminating between PsP and TTP. We sequentially added these ranked features to form a feature set and evaluated their classification performance by 10-fold CV (**Fig. 2c**). We determined the 43 top-ranked features with best classification accuracy as the optimal feature set, whose discriminative ability were further confirmed by hierarchical clustering analysis (**Fig. 2d**). The sample numbers 1 to 23 and 24 to 84 represent the PsP and TTP cases, respectively. The hierarchical clustering analysis shows a significant difference between the two phenotypes.

### Identification of the significant genes

**Figure 3.**
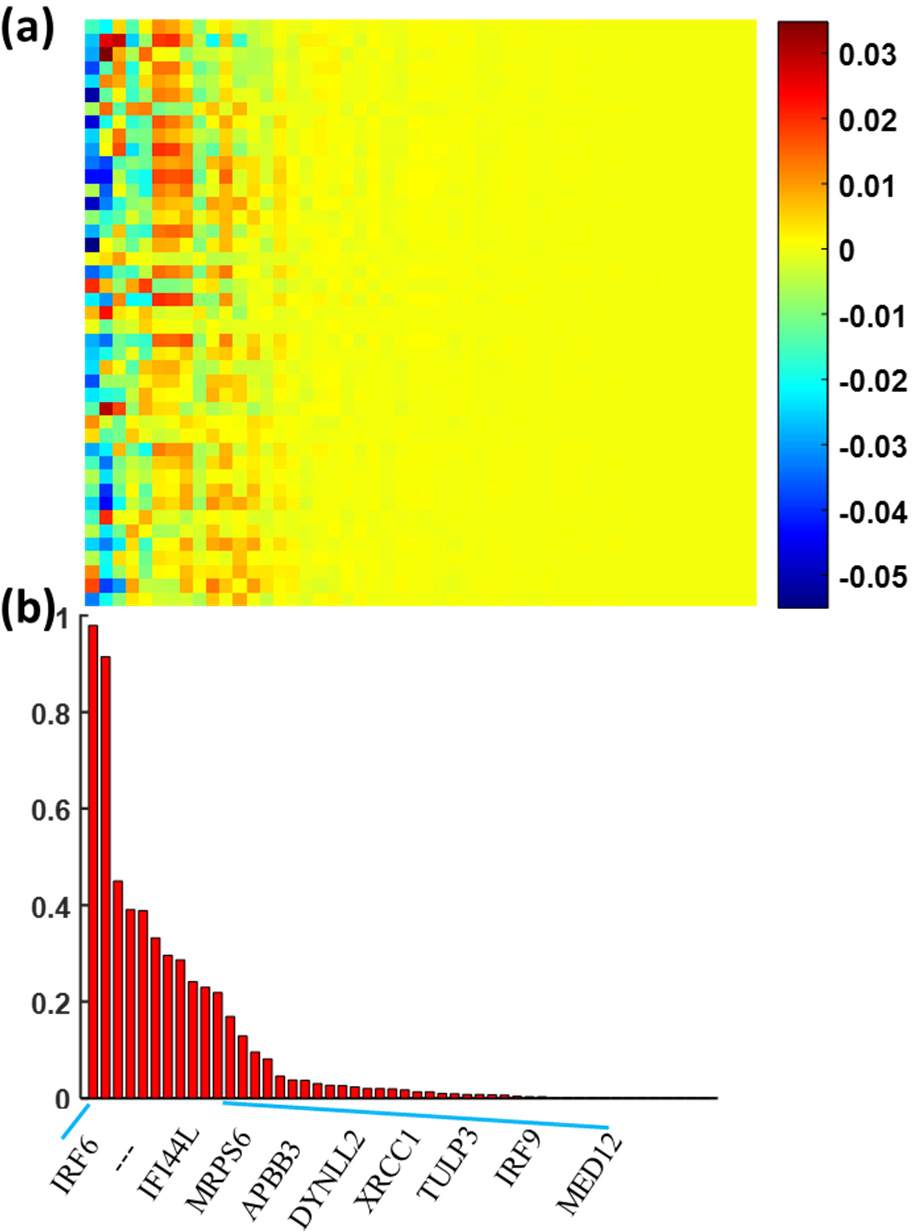
The association of the imaging features with the differentially expressed genes. Panel a shows the overall weight map of 119 genes relative to 43 imaging features and panel b shows the genes with top-ranked weights across imaging features.

In our previous radiogenomics study, we identified 119 differentially expressed genes with *p* < 0.005 using the Wilcoxon rank sum test (**Supplementary Table S1)**. We applied the sparse regression model to assess the relationship between the 119 genes and 43 imaging features from DTI. **Fig. 3a** shows the association map between the genes and imaging features. We calculated the overall weights of each gene relative to 43 imaging features and sorted them from left to right in **Fig. 3**. Twenty-three genes with top-ranked weight sum were identified as the significant genes (**Supplementary Table S2)**, which were highly correlated with the imaging features. The data from **Fig. 2c** and **d** demonstrated the discriminative ability of 43 top-ranked features.

To confirm the biological and clinical relevance of our identified significant genes, we conducted functional analysis on these 23 genes by checking their enriched signaling pathways and disease relevance. These genes are most enriched in the interferon and DNA break repair related pathways, indicating their extensive participation in the inflammation and cell cycle processes. These results further confirmed the radiogenomics method is promising in identifying the significant with high clinical relevance. More details are provided in the **Supplementary Materials**.

### Survival analysis of 2YS model on Wake dataset and TCGA dataset

**Figure 4.**
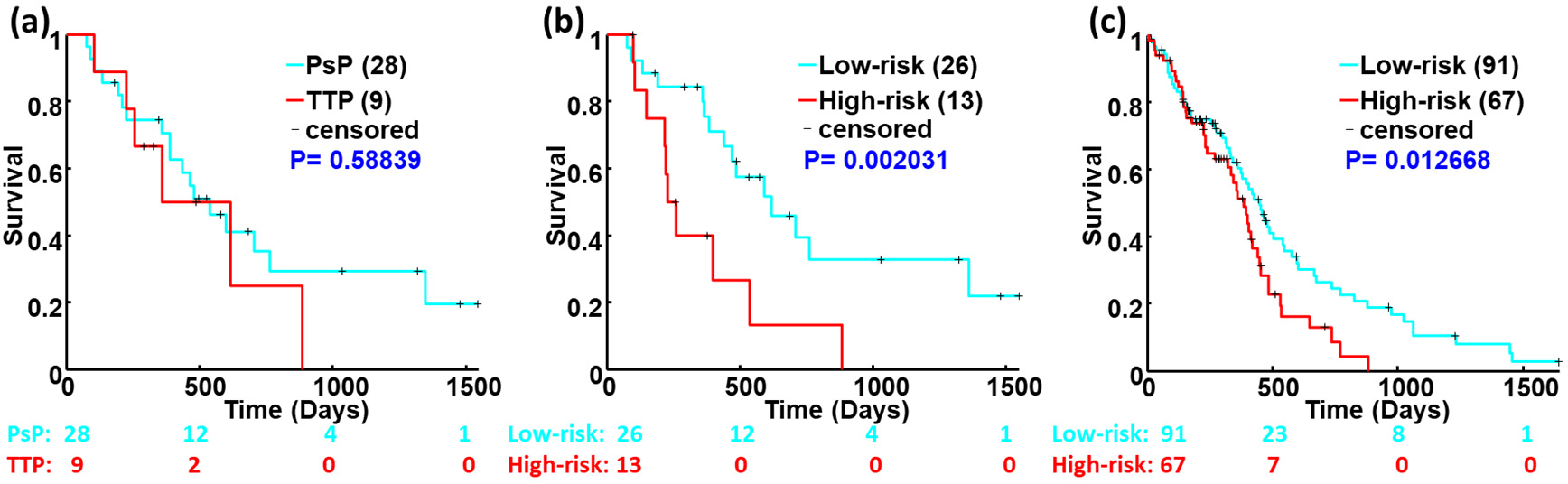
Kaplan-Meier survival analysis based on the clinical diagnosis (a) and 2YS scores using Wake dataset ((b), n=39) and TCGA dataset ((c), n=158).

We first applied the Wake dataset of 39 cases as the training data to build the 2YS prognostic model for prediction of the binary outcomes. The patients were divided into high-and low-risk groups based on their 2YS scores. The Kaplan-Meier survival curves in **Fig. 4(b)** show the significant survival difference between two groups (Log-rank *P* = 0.0020) using the Wake dataset. We then tested this model using the TCGA dataset of 158 cases, which was not used in constructing the prognostic model. Survival difference between high-and low-risk groups were statistically significant with Log-rank *P* = 0.0127, as shown in the **Fig. 4(c)**. the Kaplan-Meier survival curves reveal the consistent outcomes of 2YS predictive model on the two different cohorts, the results from TGCA data set was consistent with that from the Wake cohort. 2YS scores-yield classification results were significantly associated with the overall survival of patients, indicating that 2YS model can be used as an objective and quantitative tool for GBM diagnosis in clinical practice. Notably, The PsP and TTP groups classified based on the clinical criteria showed no significant association with survival (Log-rank *P* = 0.5884, **Fig. 4(a)**). It may be due to the variability of the grading process by individual physicians based on follow-up imaging. Since two patients in Wake dataset do not have diagnosis information of PsP or TTP in clinical records, we excluded these 2 cases and used the rest of 37 cases for survival analysis based on the criteria of clinical practice (**Fig. 4(a)**).

To further verify the effectiveness of our model, we trained and validated the model using 158 patients’ data from TCGA. TCGA data was randomly divided into two equal subsets. One subset was used to build 2YS model and another set to validate it. Each subset was running twice as testing and validation samples, respectively. As shown in **Fig. 5**, the low-and high-risk groups from the training dataset or the validation dataset display a significant survival difference (Log-rank *P*<0.01). These results confirmed the predictive effectiveness of our proposed 2YS prognostic model.

**Figure 5.**
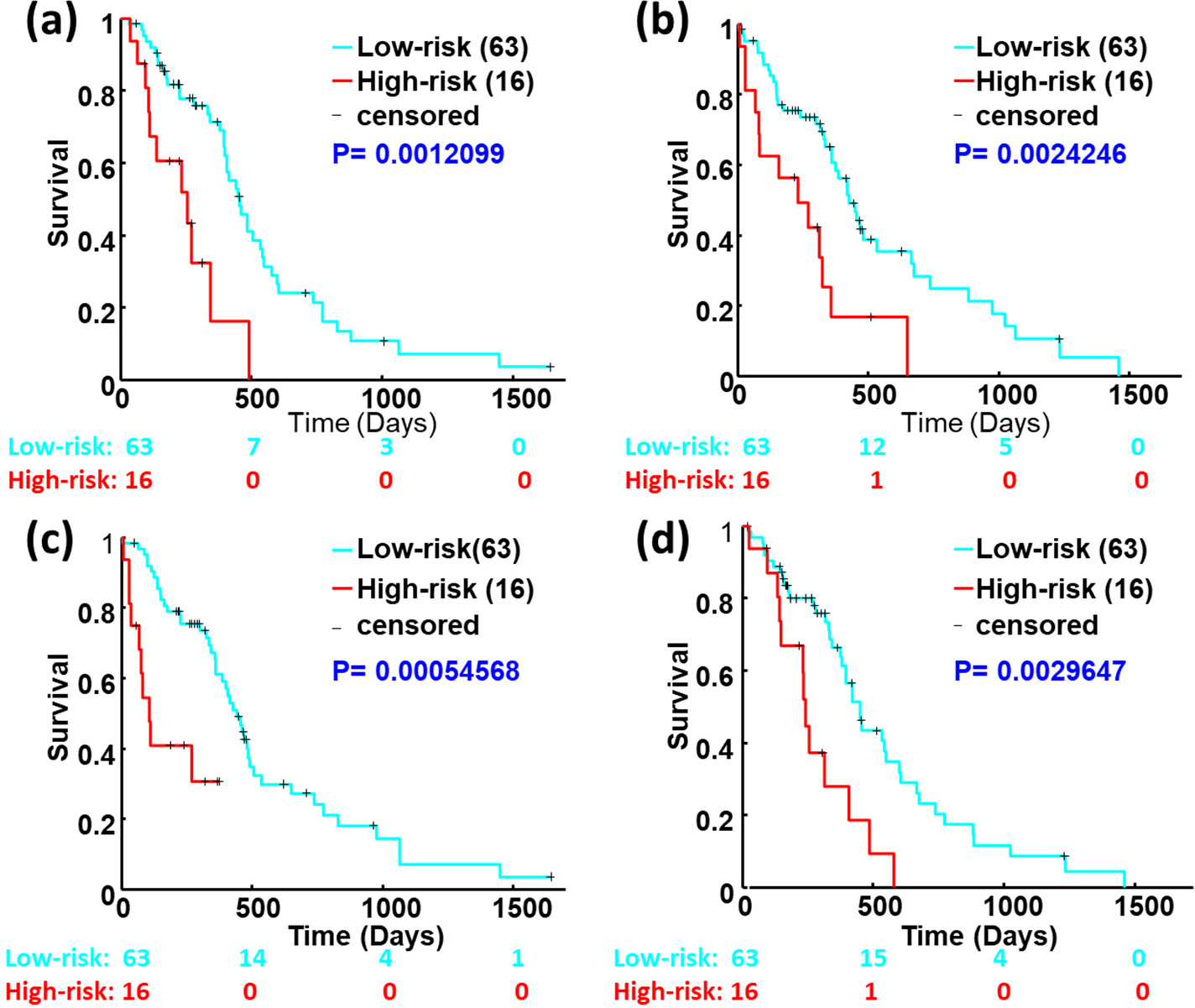
Kaplan-Meier survival analysis of the training and validation data sets using the TCGA-based 2YS model. 158 cases from TCGA were randomly divided into two equal subsets. Each subset was running twice as a testing and validation sample. Panel a and b show survival analysis results from the subset A as training and validation samples. Panel c and d show survival analysis results from the subset B as training and validation samples, respectively.

### Relevance between clinical characteristics and 2YS score

To reveal the correlation between clinical factors and 2YS score, we first used the 2YS score to classify the patients into low-and high-risk groups for Wake and TGCA cohorts and then performed the Kolmogorov-Simonov test on continuous variables and Fisher exact test on discrete variable between two groups, as shown in **Table 1**. We investigated the association of clinical factors of Wake dataset with patients’ survival, including age, gender, race, resection type and MGMT. Only resection type was slightly associated with 2YS scores (Log-rank *P* < 0.03). We also analyzed the clinical variables of TGCA dataset, including age, gender, race, resection type, ethnicity, KPS, IDH1, MGMT, normal cells ratio and tumor cells ratio. IDH1 mutation status was significantly associated with the risk groups (Log-rank *P*-value of 0.0001). Previous studies have shown that IDH1 mutation was a potential biomarker for GBM^36,37^. For example, Moteqi et al. found that IDH1 mutation was associated with PsP and TTP^14^.

**Table 1.**
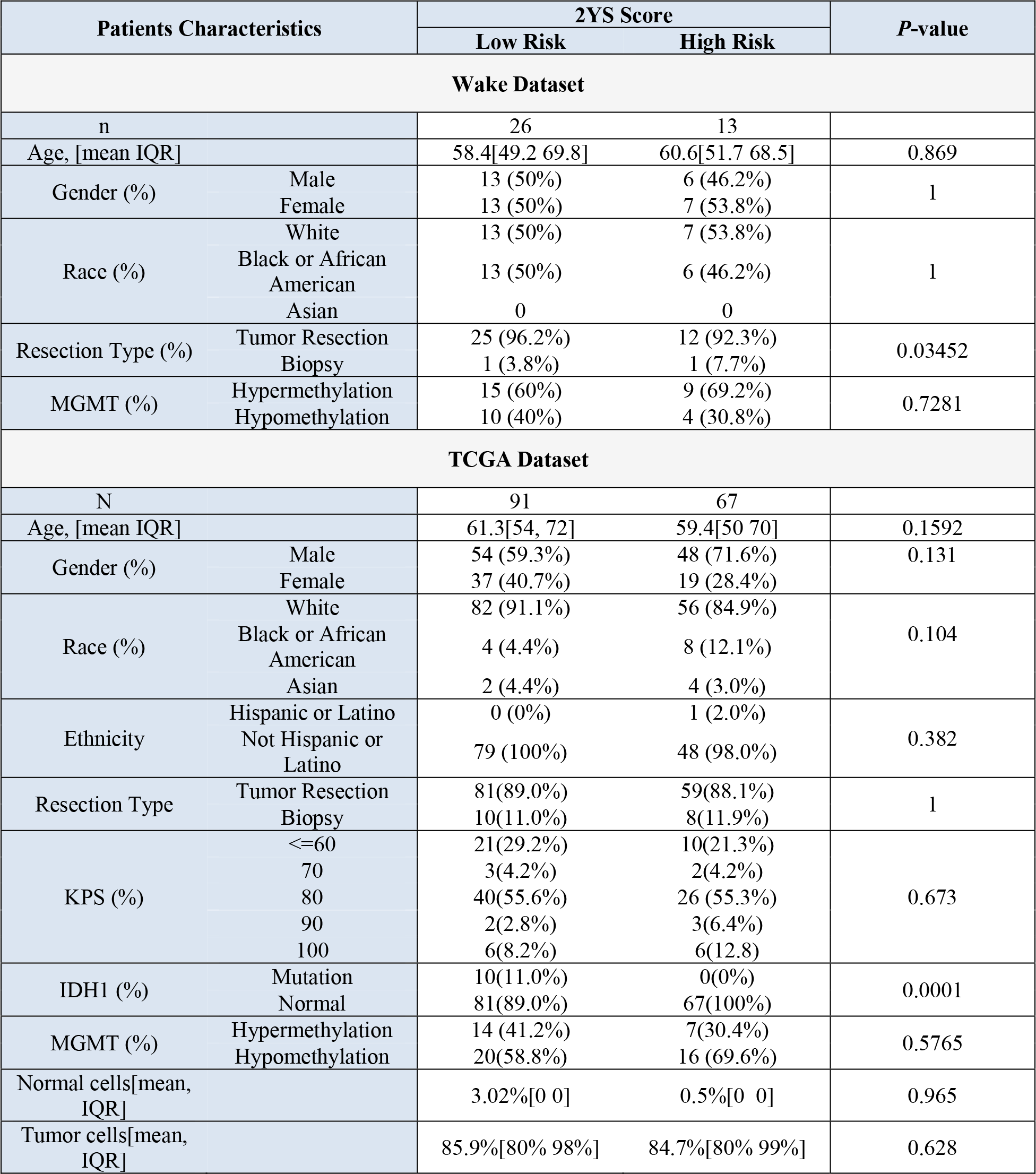
Patient characteristics categorized by 2YS score.

We assessed the prognostic performance of the 2YS score in the context of other known prognostic factors using the multivariate Cox proportional hazards model, as shown in **Table 2**. The 2YS score yielded from the Wake dataset was significantly associated with patient survival (**Table 2(a)**). We used multivariate Cox proportional hazards model to test whether individual clinical factors (gender, age, race, surgery type, and MGMT) from Wake dataset could predict patients’ survival and found that their predictive ability was low. We also constructed multivariate Cox proportional hazards model using TCGA data set (**Table 2(b**)). The 2YS score from this model was significantly associated with the overall survival, while most of other clinical factors, including gender, age, KPS, normal cell ratio, were not or marginally correlated with patients’ survival. Tumor cell ratio was the only clinical factor significantly associated with patients’ survival (p-value = 0.038).

**Table 2.**
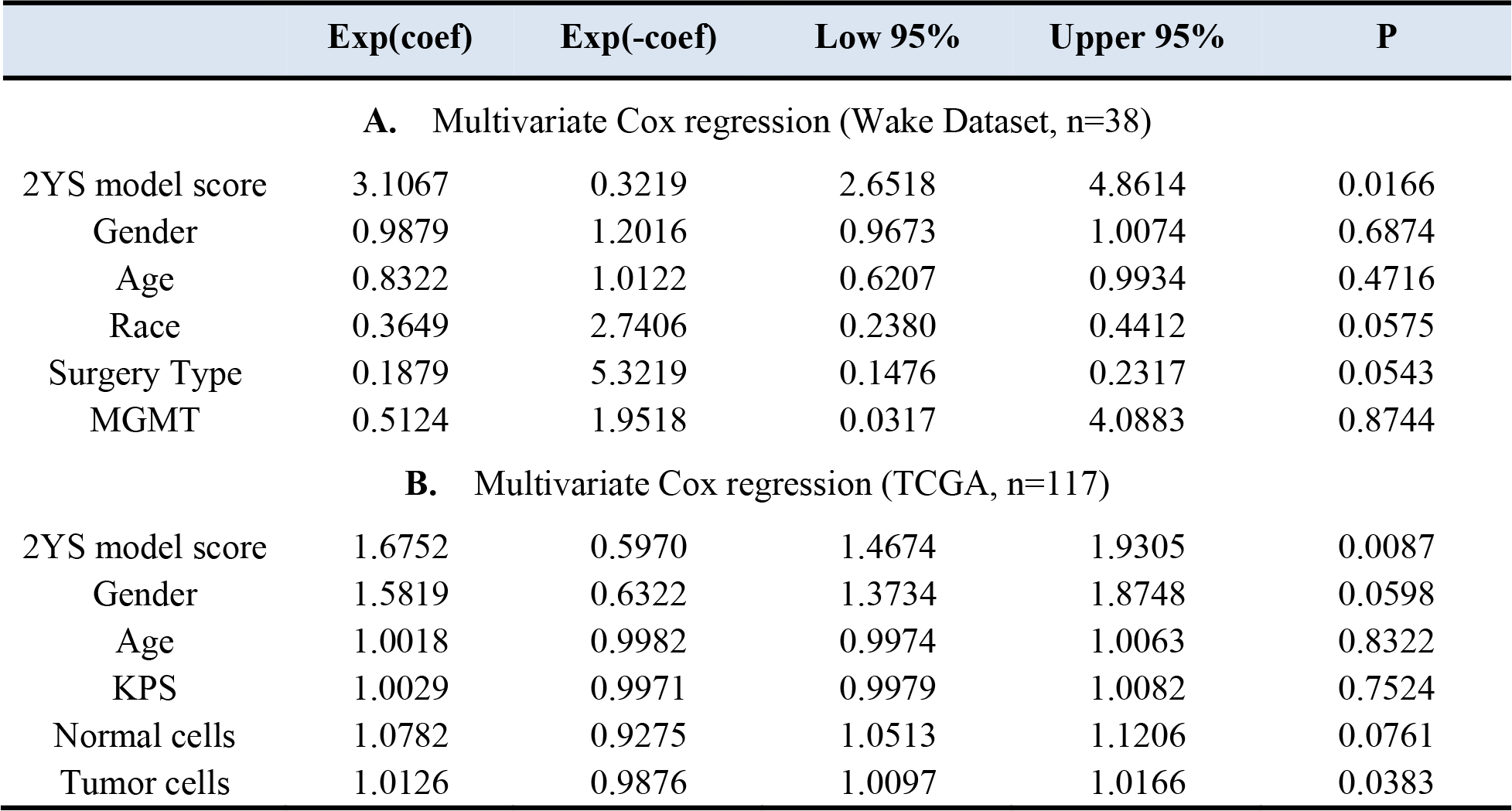
Multivariate Cox Proportional Hazards Model for Overall Survival

### Assessing significance of selected genes

To evaluate the significance of gene features in the 2YS model, we performed a bootstrap analysis with 1000 bootstrap iterations on TCGA dataset. The 95% CIs of the coefficient estimates of the 2YS model for each gene’s expression value were yielded. We identified 12 out of 23 features as the significant features in the 2YS model, as shown in **Table 3**. Most of these genes have been previously reported to participate in critical biological processes related to cell cycle and DNA repair. In particular, IRF9 is involved in a series of biological pathways directly or indirectly related to cell proliferation, apoptosis, and innate immunity^38–44^, and has been identified as a potential biomarker for distinguishing PsP and TTP in our previous radiogenomics study^35^. Functional analyses of these genes (**Supplementary materials**) indicated that IRF9, MGLL and OAS1/3 interact with IFNβ and interferon alpha to exert inflammation functions; while XRCC1, TCTN2 and FOXJ2 act synergistically with TP53 and ADAMTS8 in the regulation of cell cycle-related processes.

**Table 3:** Identification of significant gene features in 2YS model by bootstrap analysis.

|  | Feature (genes) | 95% low | 95% high |
| --- | --- | --- | --- |
| 1 | IRF9 | 0.5721 | 0.8568 |
| 2 | OAS3 | -0.9602 | -0.3276 |
| 3 | TULP3 | -0.8305 | -0.3923 |
| 4 | IFI6 | -0.6036 | -0.0959 |
| 5 | MGLL | 0.4811 | 1.4234 |
| 6 | BAZ2A | 0.2777 | 0.6136 |
| 7 | ERCC1 | -0.9589 | -0.0290 |
| 8 | OAS1 | 0.0451 | 0.7442 |
| 9 | FOXJ2 | -0.7236 | -0.0889 |
| 10 | RAB8A | -0.9366 | -0.2549 |
| 11 | TAB3 | 0.1637 | 0.8704 |
| 12 | TCTN2 | 0.0467 | 0.8706 |

## Discussion

In this study, we aimed to develop a prognostic model to evaluate tumor progression potential of GBM patients following standard therapies. To achieve this purpose, we applied a dictionary learning scheme to obtain imaging features of GBM patients with PsP or TTP from the Wake dataset. Based on these radiographic features, we then conducted radiogenomics analysis to identify the significantly associated genes. These significant genes were then used as features to construct a 2YS logistic regression model. GBM patients were classified into low-and high-survival risk groups based on the individual 2YS scores derived from this model. We tested our model using an independent TCGA dataset and found that 2YS scores were significantly associated with the patients’ overall survival. To further verify the effectiveness of our model, the TCGA dataset were randomly divided into 2 cohort. Each cohort were split into training and validating groups. Our results show that the training and validation datasets displayed a significant survival difference (Log-rank *P*<0.01), confirmed the effectiveness and robustness of our proposed 2YS model in predicting clinical outcomes of GBM patients after standard therapies. However, most of other clinical factors, including gender, age, KPS, normal cell ratio, were not or marginally correlated with patients’ survival.

Most of the existing work involved in quantitative imaging have required interest of area (ROI) determination of lesions through the laborious outline, semi-/automated segmentation by algorithms, followed by the measurement of expert predefined features, such as size, color, shape, and texture, for describing lesions characteristic. In the case of our work, the ROI determination of GBM on DTI is a tough task due to the low imaging quality^31^, although DTI has the potential to differentiate the tumors and brain tissues with treatment effects^29,30^. After training of the dictionary learning model using expert-derived label annotations, our feature extraction and selection were achieved without relying on segmentation and manual steps, which greatly increases its scalability. Additionally, the conventional predefined descriptors may be unqualified for completely characterizing the tumors. In contrast, our previous developed classification-oriented dictionary learning scheme provides the specific and common features for different treatment results, enabling identification of imaging features related to the clinical treatment progression^31^.

In recent years, radiogenomics emerges as a new research field for discovering the relationship between imaging and genetics profiles of patients, with the purpose of better understanding and promoting the research for cancer treatment^25^. For example, the association map, determined by the radiogenomics study, can discover the specific genetics underlying the imaging characteristics^45^ and construct the predictive radiographic signatures^46^. In this study, we conducted the radiogenomics study to explore the relationship between DTI phenotype and genotype (i.e., gene expression) for determination of significant genes with our previously developed low-rank sparse regression model. Through this radiogenomics study, these identified significant genes were confirmed to reflect the development of PsP and TTP well, since they were highly associated with the imaging features, and these imaging features from classification-oriented dictionary learning method could provide a good classification performance for PsP and TTP (**Fig. 2**).

Actually, we have further documented the clinical relevance of these significant genes by a series of subsequent functional analysis. Specifically, after obtaining the 23 significant genes from the radiogenomics study, the IPA (Ingenuity Pathway Analysis) software was employed to explore the biological functions of these genes, including critical signaling pathways they participate, diseases or functions they are associated with, and biological network they are involved. The canonical pathway analysis tool was used to determine the most significantly affected pathways in the context of established signaling and metabolic pathways, determine which pathways overlap based on molecules in common. Enrichment significance and candidate gene coverage profile are calculated. The disease/function annotation tool provides details associated with the disease or biological function such as related molecules, known drug targets. The network analysis tool was applied to build and explore transcriptional networks, microRNA-mRNA target networks, phosphorylation cascades and protein-protein or protein-DNA interaction networks. As shown in the supplementary materials, **Fig. S1** illustrates the top-ranked signaling pathways associated with the 23 candidate genes. IRF9 is involved in the interferon signaling pathway, while XRCC1 takes part in both BER (base excision repair) and DDSB (DNA double-strand break) repair pathways. **Table S3** lists the most relevant (P<10^−3^) diseases and biological functions these genes are implicated. IRF9 and MGLL are involved in inflammation process; ERCC1 and XRCC1 are associated with cell cycle (G2/M) process. **Fig. S4** shows the interaction network between the significant genes and other related molecules. IRF9 and OAS1/3 interact with inflammatory genes IFNb and interferon alpha, and also indirectly associates with histone 3 and 4; while XRCC1, TCTN2, and FOXJ2 synergistically interact with TP53 and ADAMTS8 to exert their cell cycle related functions.

Despite annotated merits, radiogenomics approach has the other side of a coin, i.e., integrating of phenotype and genotype data from each patient objectively and inevitably limited the sample size for significant genes determination, which is also a probably imperfection of this study. It is a challenge task to simultaneously collect all types of datasets from individual patients, including DTI, gene expression, and clinical records, because these datasets were generated for clinical purpose and not specially designed for academic research. Also, TCGA and the Cancer Imaging Archive (TCIA) do not include the clinical records about treatment results (i.e., triage of patients with standard therapies) and DTI; thus, TCGA dataset in this study could not be used to perform radiogenomics study for exploring the significant gense, but it can be employed for validation of the 2YS model. However, owing to the successful identification of significant genes in Wake dataset, we have reached a good result from a small set of training samples and independently validated the model on TCGA dataset with desirable performance. This suggests that we have derived a powerful prognostic model. Before translation of this system for use in clinical practice, this model should be trained on the dataset from diverse cohorts and tested on additional independent cohorts to evaluate the model’s generalizability better.

**Fig. 4(a)** presents an unexpected and intriguing result that there is no significant survival difference with 0.5884 of Log-rank *P* value between PsP and TTP groups by clinical diagnosis from Wake dataset. This result is unsurprised since it is not an isolated case. For example, recently several work uncovered no significant association with survival between different stages of breast cancer on some cohorts by current clinical criterion^26,47^. This motivated us to explore a novel prognostic marker for stratification of GBM patients into different risk groups with significant survival association. There are several potential factors caused this abnormal phenomenon (**Fig. 4(a)**). First, the clinical practice is to perform the follow-up MRI examinations based on the morphological changes of lesions. It usually takes several months to make the final diagnosis and causes an adverse effect on the clinical management of patients. Second, the clinical diagnosis criterion for PsP and TTP is defined on the phenotype progression, which is unable to recover the diseases essential progression comprehensively. Pathological confirmation presents the reliable diagnosis, whereas brain tumor biopsies are not usually applied in clinical practice due to its invasiveness, increasing the risk for subsequent therapy. Thus, the imaging-based diagnosis may lead to the insignificant survival difference between two entities. Finally, the limited sample size may result in the outcome bias. We will further collect cases to verify this discovery, although it will take a long time. In our previous work, we have tried to overcome the critical delay of diagnosis (i.e., the first problem mentioned above) through developing a computer-aid diagnosis system based on DTI^31^. To deal with the second issue noted above, we integrated imaging features and gene expressions to determine the 23 significant genes as features for construction of a predictive survival model, which was significantly associated with survival.

From the view of clinicians, our 2YS score provides an innovative diagnosis index, which bridges the radiogenomic biomarkers and clinical survival status. Physicians can utilize the 2YS score to give reliable estimations of the GBM progression risk. Compared to current medical judgments, the 2YS score only needs partial retrospective information of patients’ genomic and radiographic records. In this way, the long-time followup processes and invasive pathologic detections are avoided. Thus, 2YS score can be considered as an actionable clinical assistance index of judging GBM prognostics. In addition, the 2YS score is verified through the statistical association with some known omics biomarkers. In **Table 1**, “IDH1 mutation” is highly associated with high & low risks of 2YS Score (P-values < 0.05). Besides, we discovered that “tumor cells” owns the similar GBM survival risk as 2YS score in **Table 2** (P-value: 0.0383). In sum, our proposed 2YS score proves to be clinical meaningful and an important diagnostic complement for GBM physicians.

## Supplemental Figures and Tables

### Figure legends

**Figure S1.**
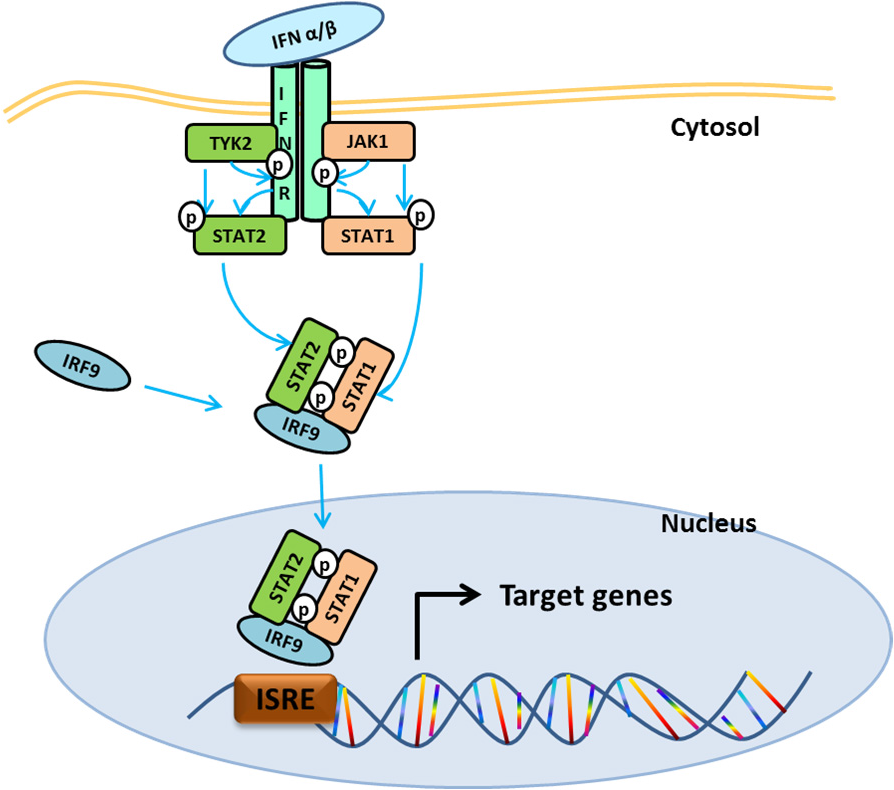
JAK-STAT pathway for type I IFN, retrieved and summarized from previous studies.

**Figure S2.**
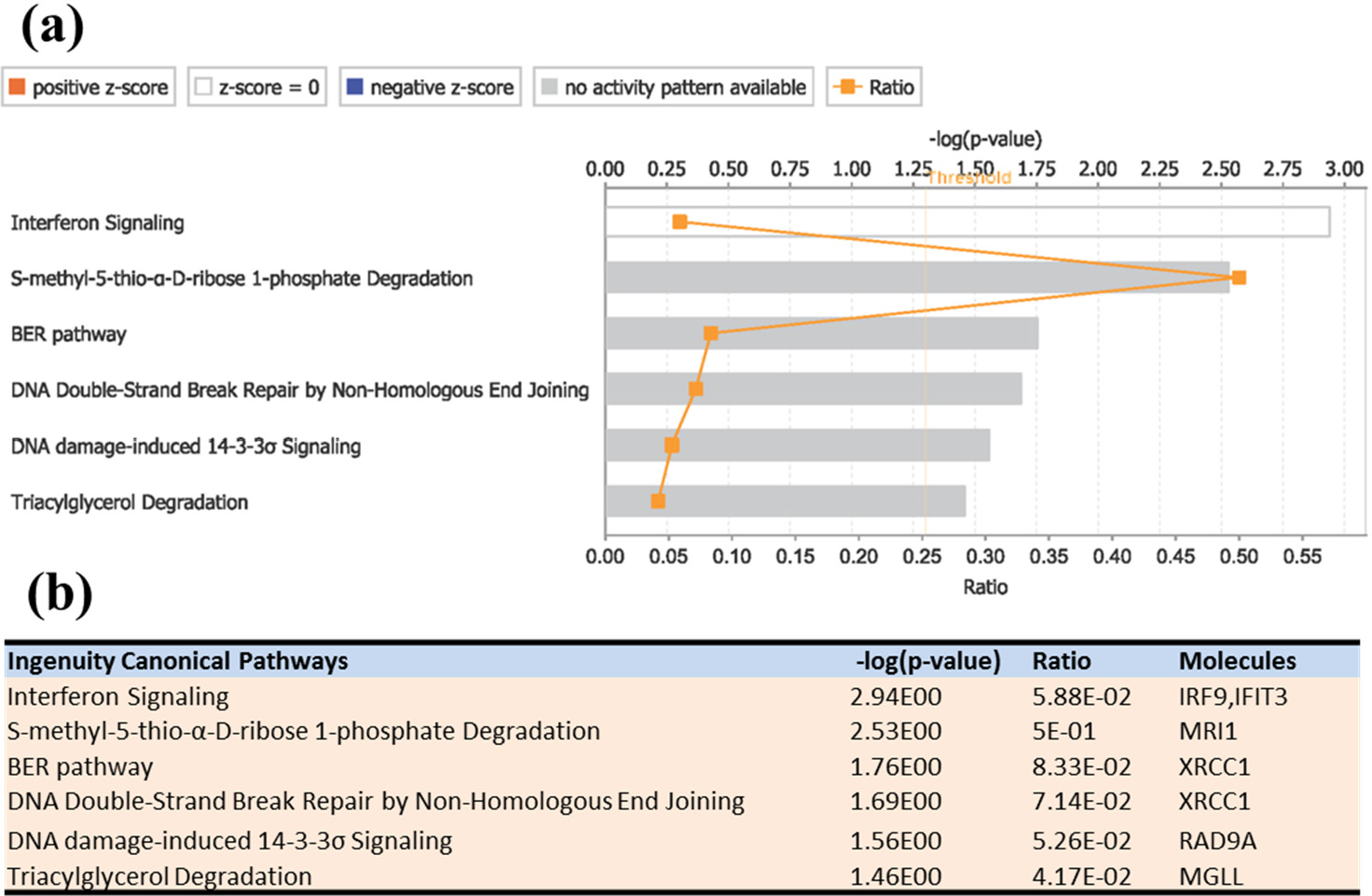
Top-ranked Canonical Pathways associated with the **<u>33 candidate genes selected by radiogenomics</u>**. Canonical pathways are ordered by the *p*-values (*p*<0.05).

**Figure S3.**
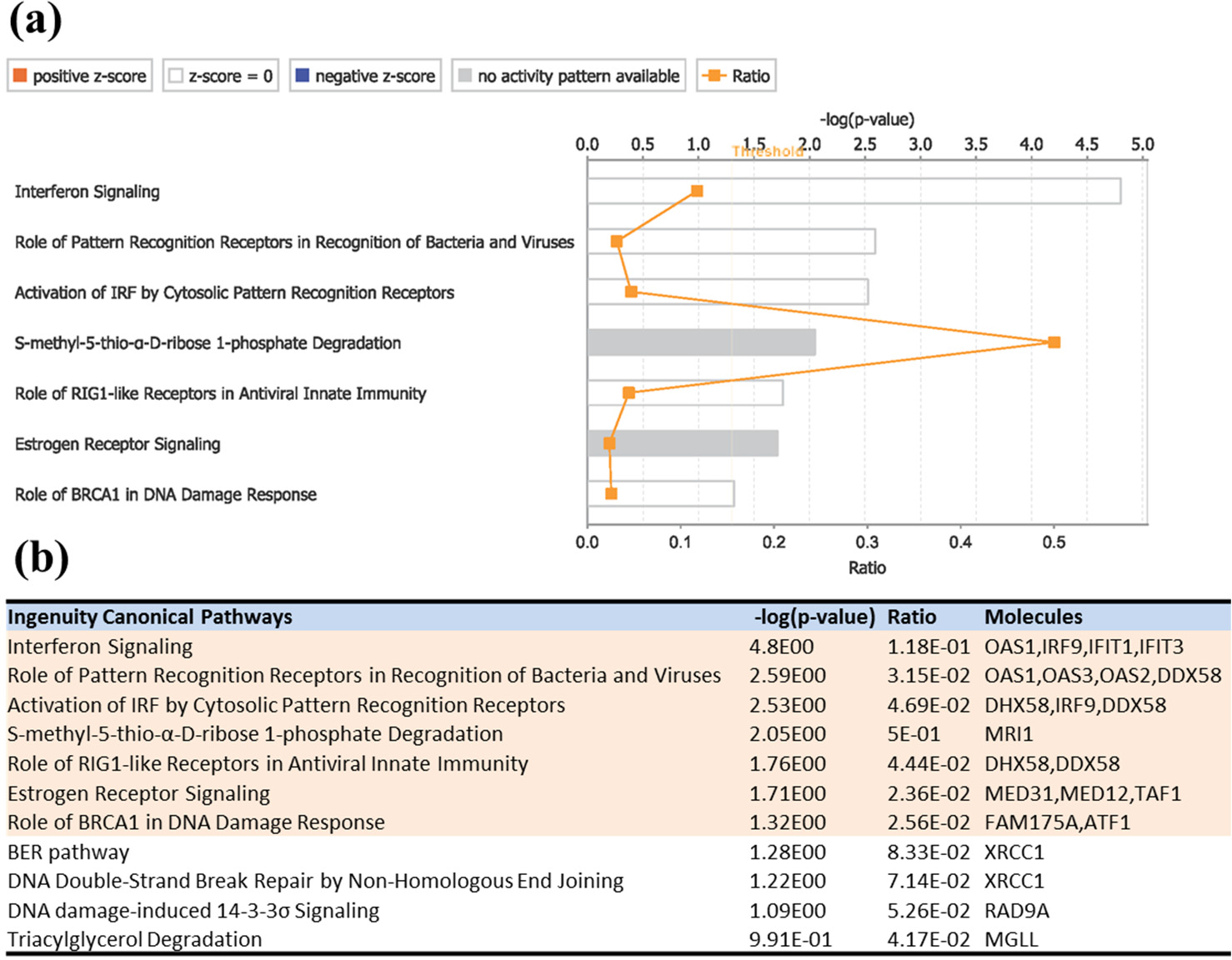
Top-ranked Canonical Pathways associated with the 119 differentially expressed genes. Top-ranked Canonical Pathways were highlighted red in (b) with p<0.05.

**Figure S4.**
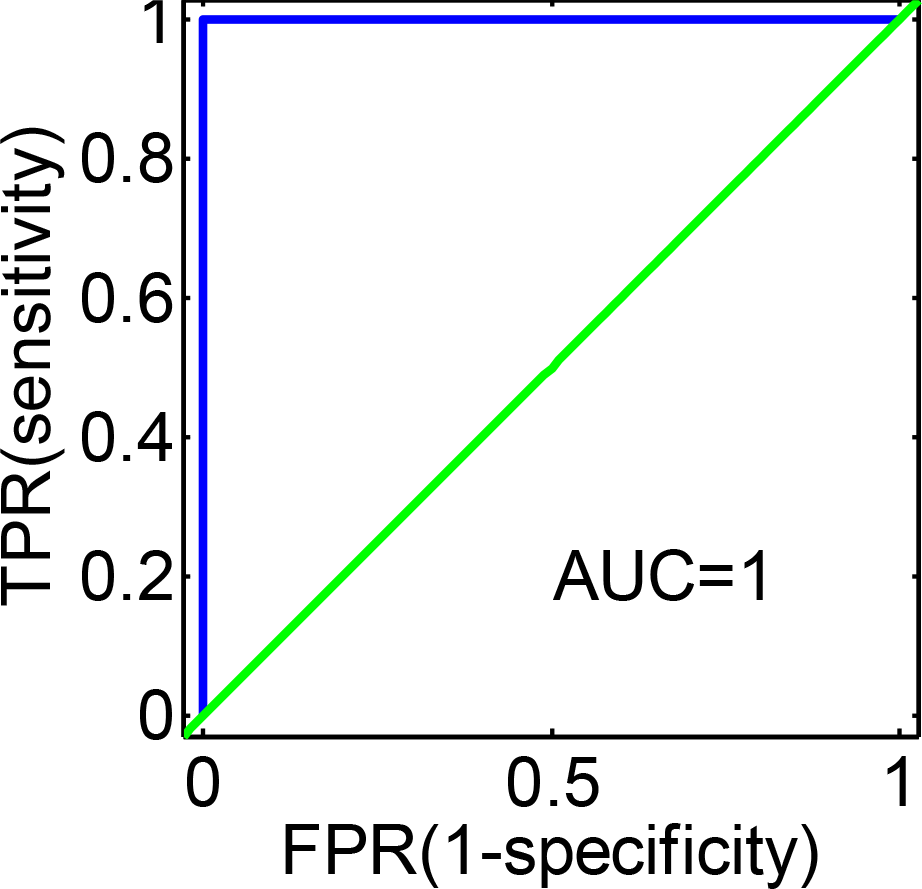
The ROC curve for the classification of the 119 differentially expressed genes as features. The SVM classifier and fivefold cross-validation were applied in this study. *sensitivity* = *TP*/(*TP* + *FN*) and *specificity* = *TN*/(*FP* + *TN*), where *TP*, *FP*, *FN*, and *TN* refer to truepositive, false positive, false negative, and true negative, respectively.

**Figure S5.**
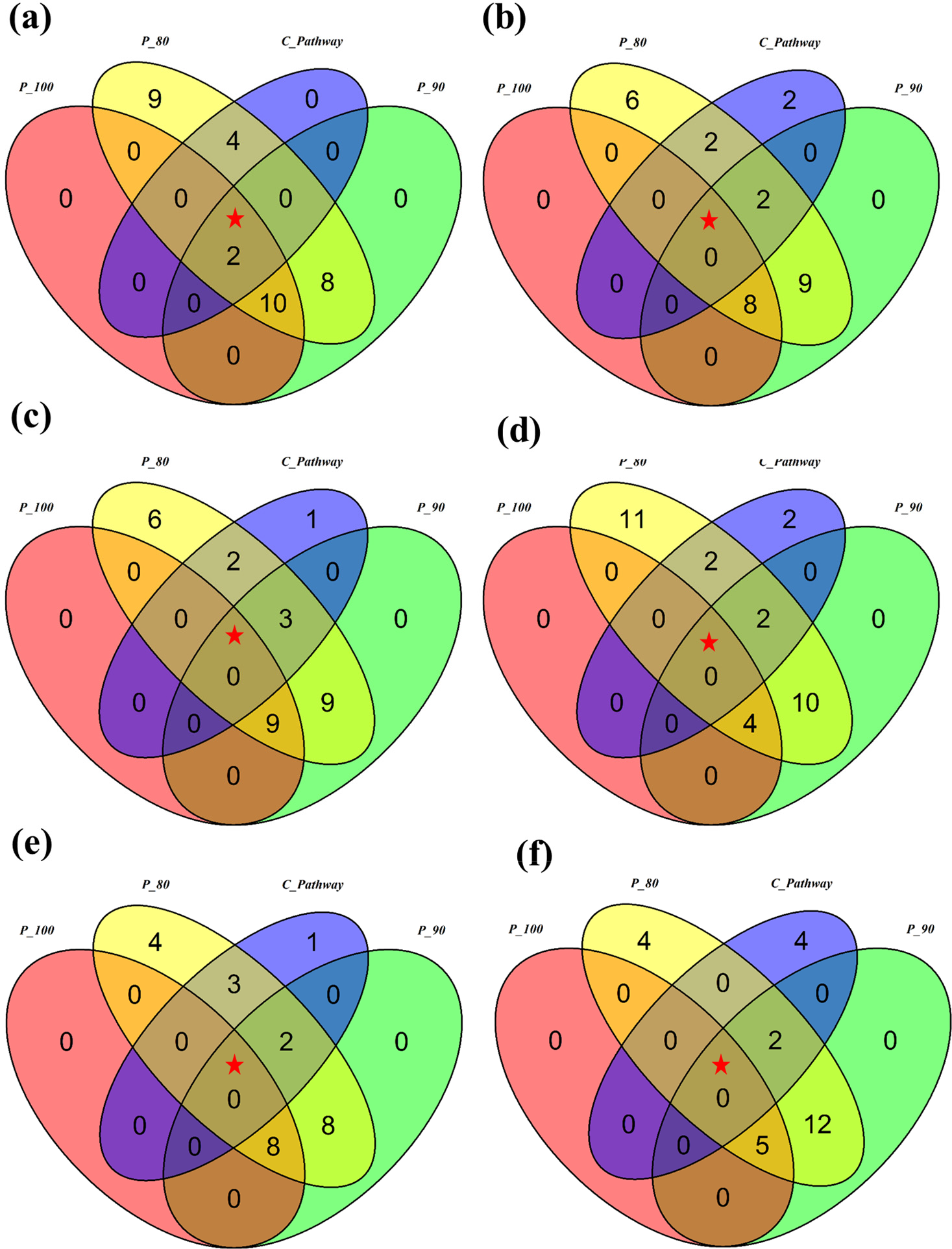
The gene overlaps among candidate genes from different coverage rates *P* and genes from top-ranked canonical pathways (i.e., *C_Pathway*). The coverage rate *P* was defined as the occurrence frequency of genes on the lists of weight ranker top 50 in 27 parameter combinations. *P_100*, *P_90* and *P_80* mean 100%, 90% and 80% of coverage *P*, respectively. Experiments with six feature sets: (a) 225 image features; (b) 50 image features used in clinic practices; (c) 50 image features randomly selected from (a); (d)-(f) 20 image features randomly selected from the 50 clinical features in (b) for three times.

**Figure S6.**
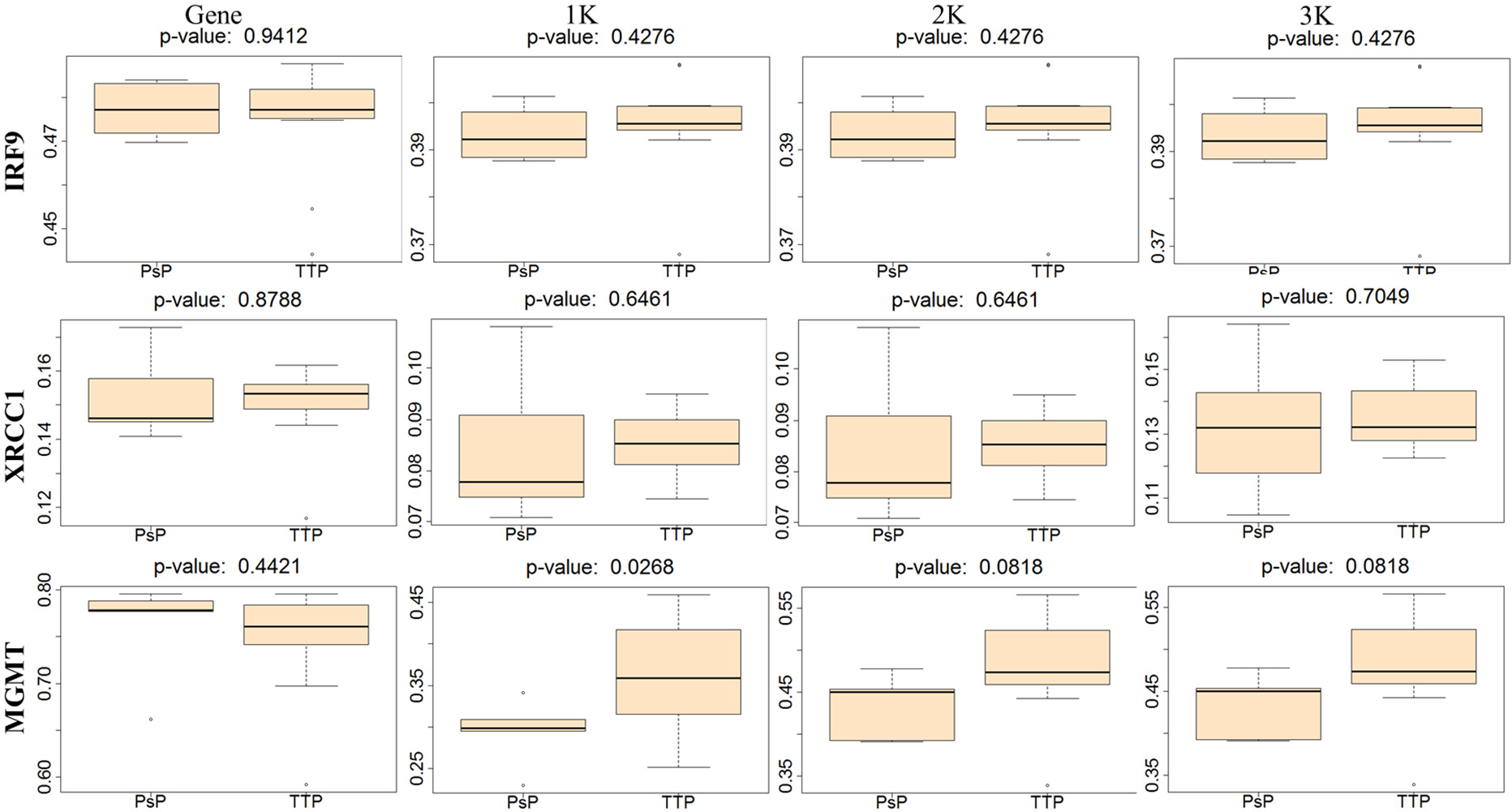
Boxplots of gene methylation, promoter methylation with 1k, 2k, and 3k window of TSS from 1 to 4 columns, respectively, for IRF9 (1^st^ row), XRCC1(2^nd^ row), and MGMT (3^rd^ row).

**Figure S7.**
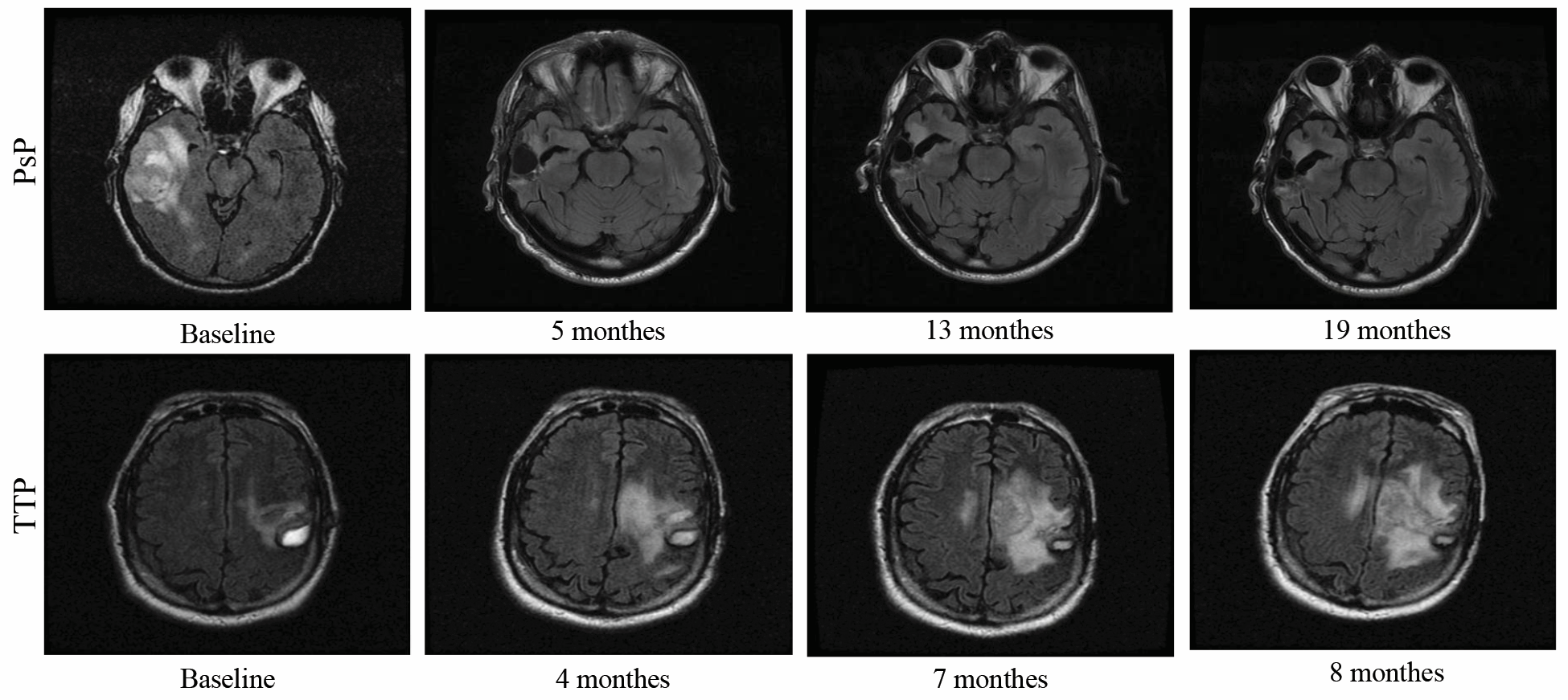
Demonstration of PsP and TTP based on longitudinal MRI. All of the images are from TCIA. The sample ID for the top row images is TCGA-06-0185, and the bottom row is TCGA-06-1084.

**Figure S8.**
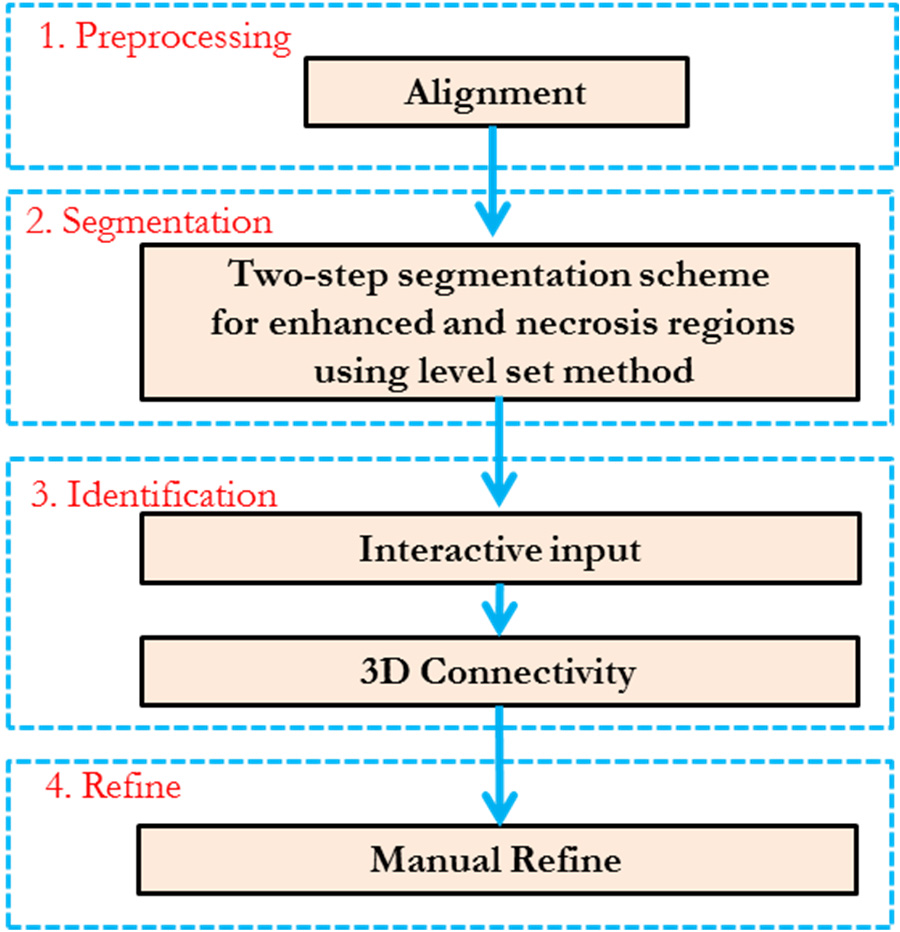
The flowchart of the segmentation of tumor regions (i.e., enhanced region and necrosis regions).

### Table legends

**Table S1.**
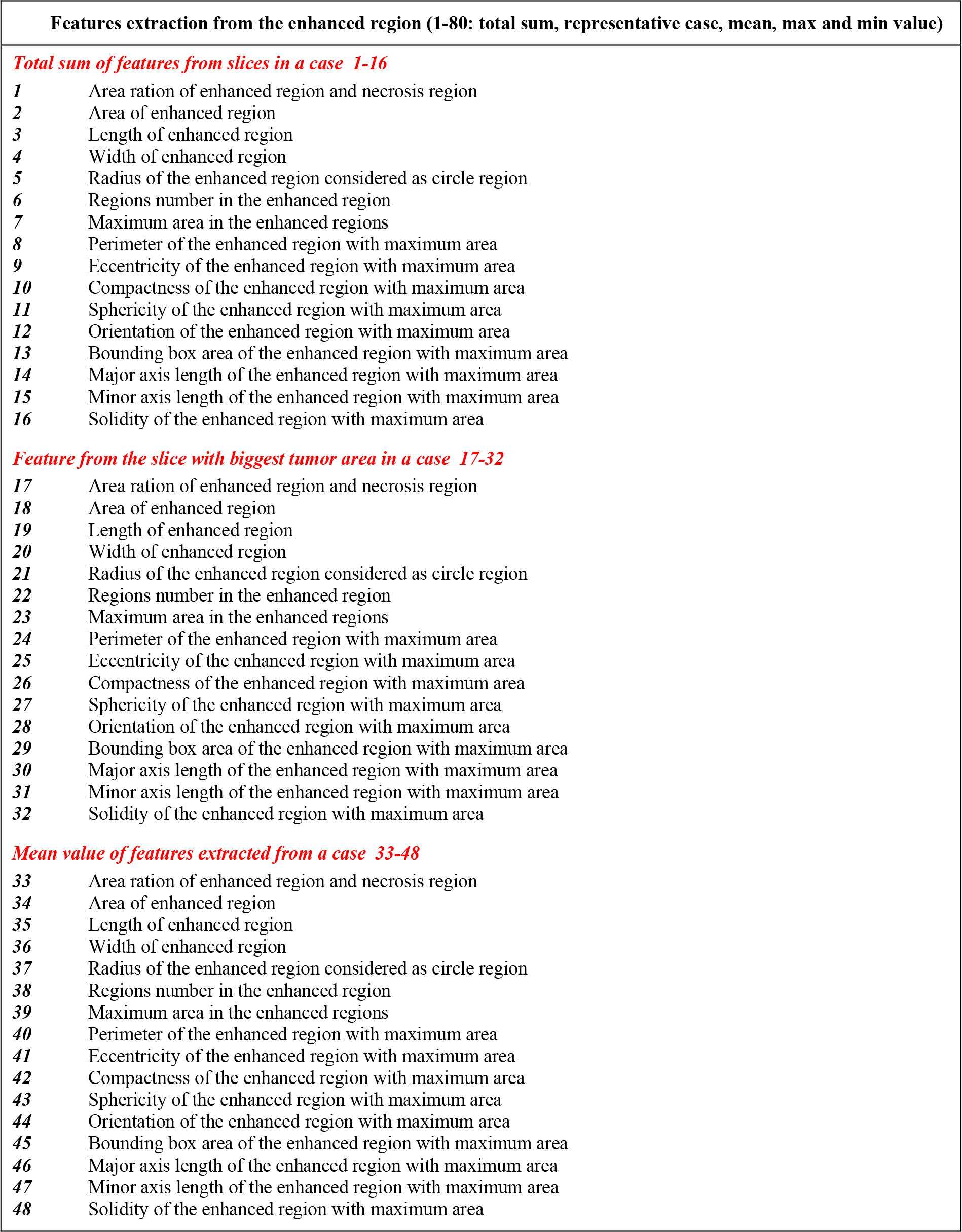
Summary of all the features extracted from the longitudinal MRI

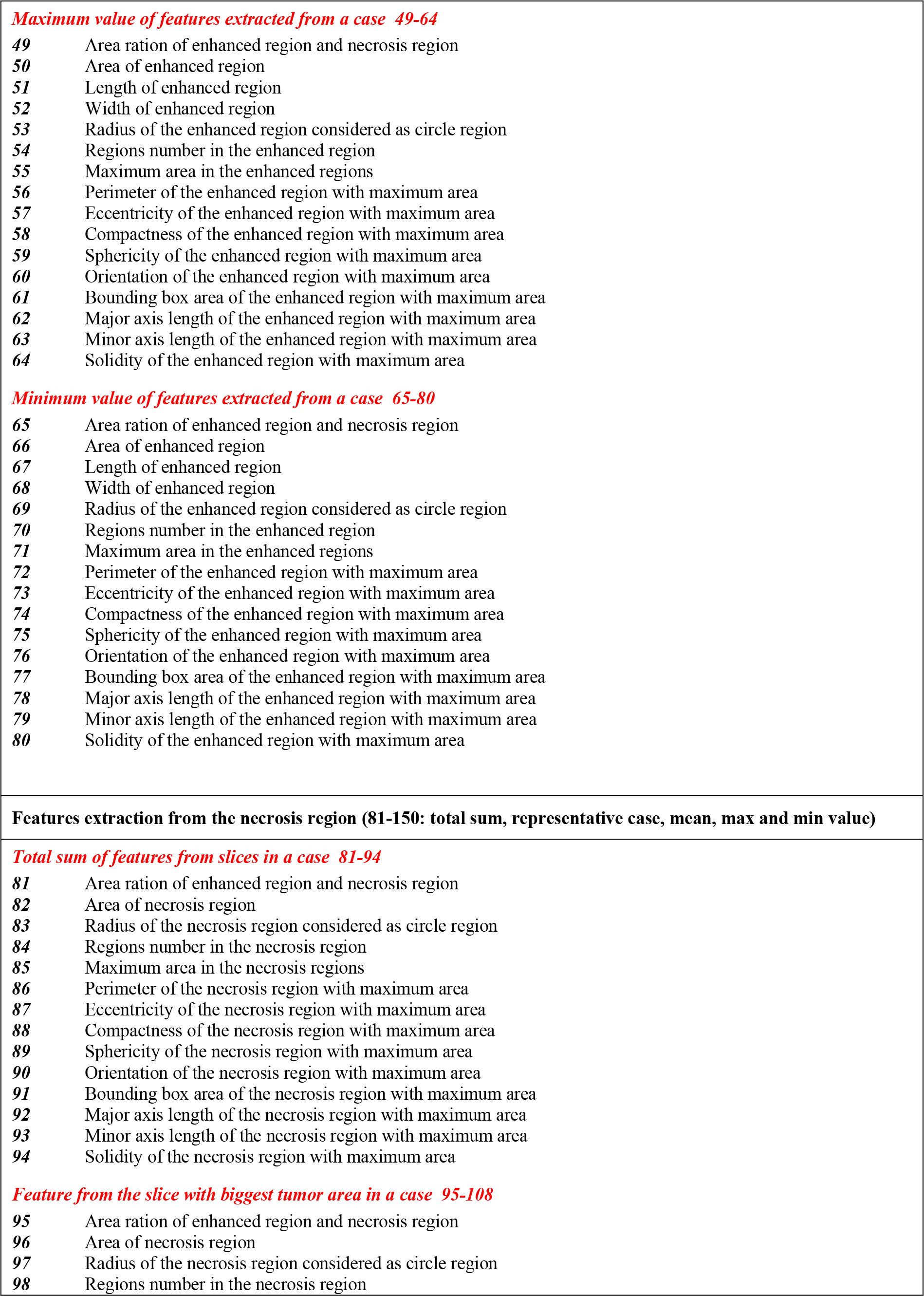

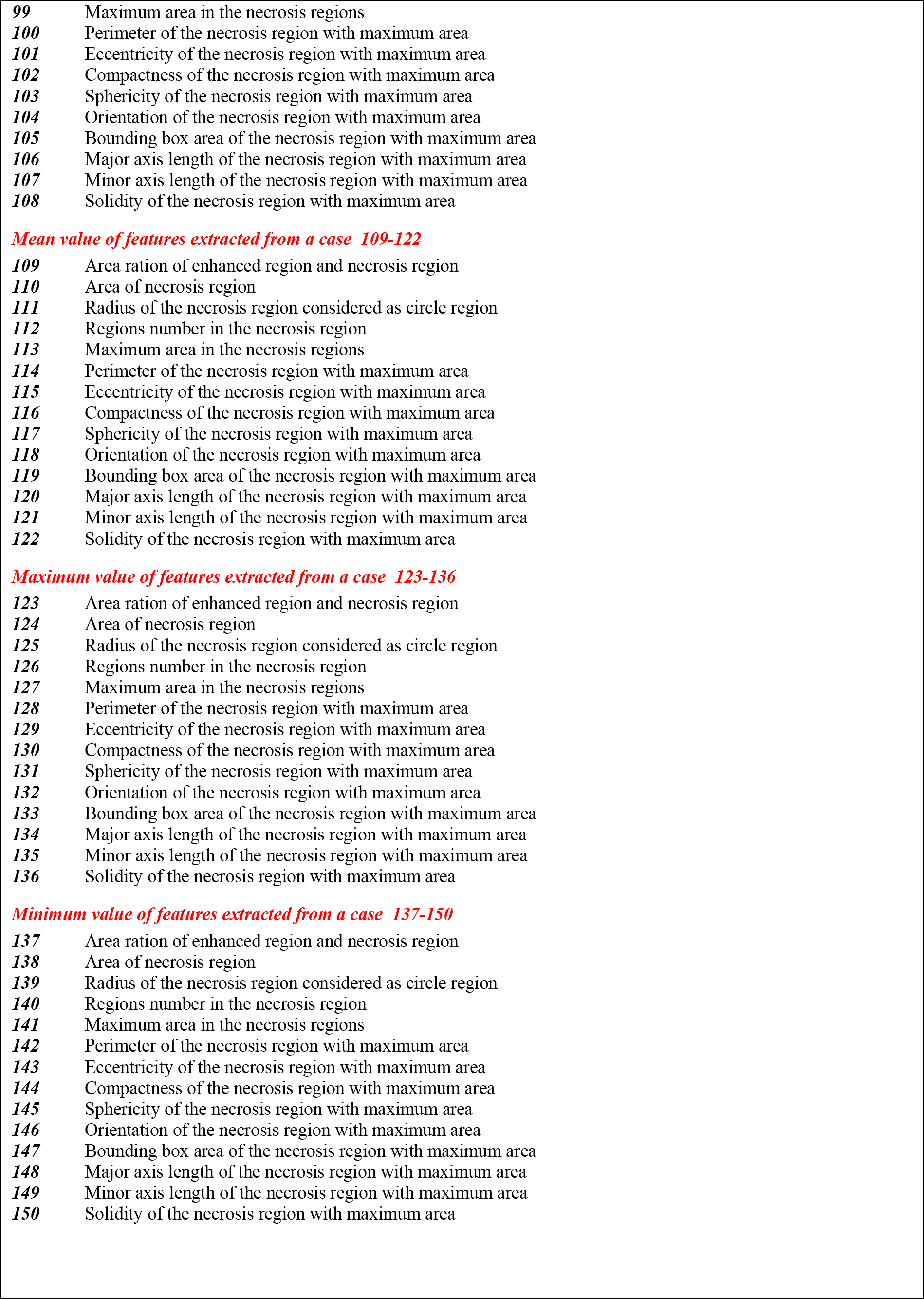

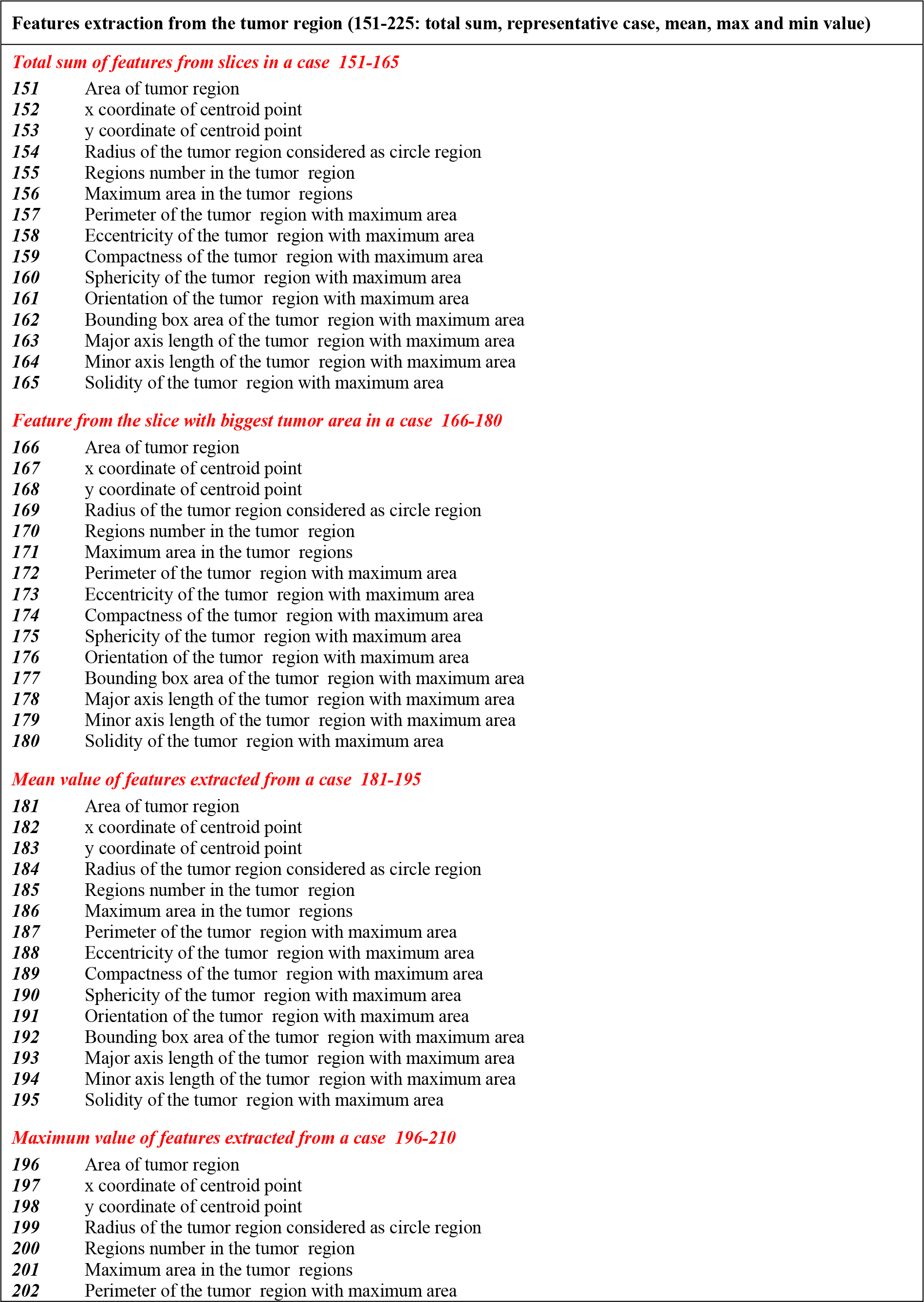

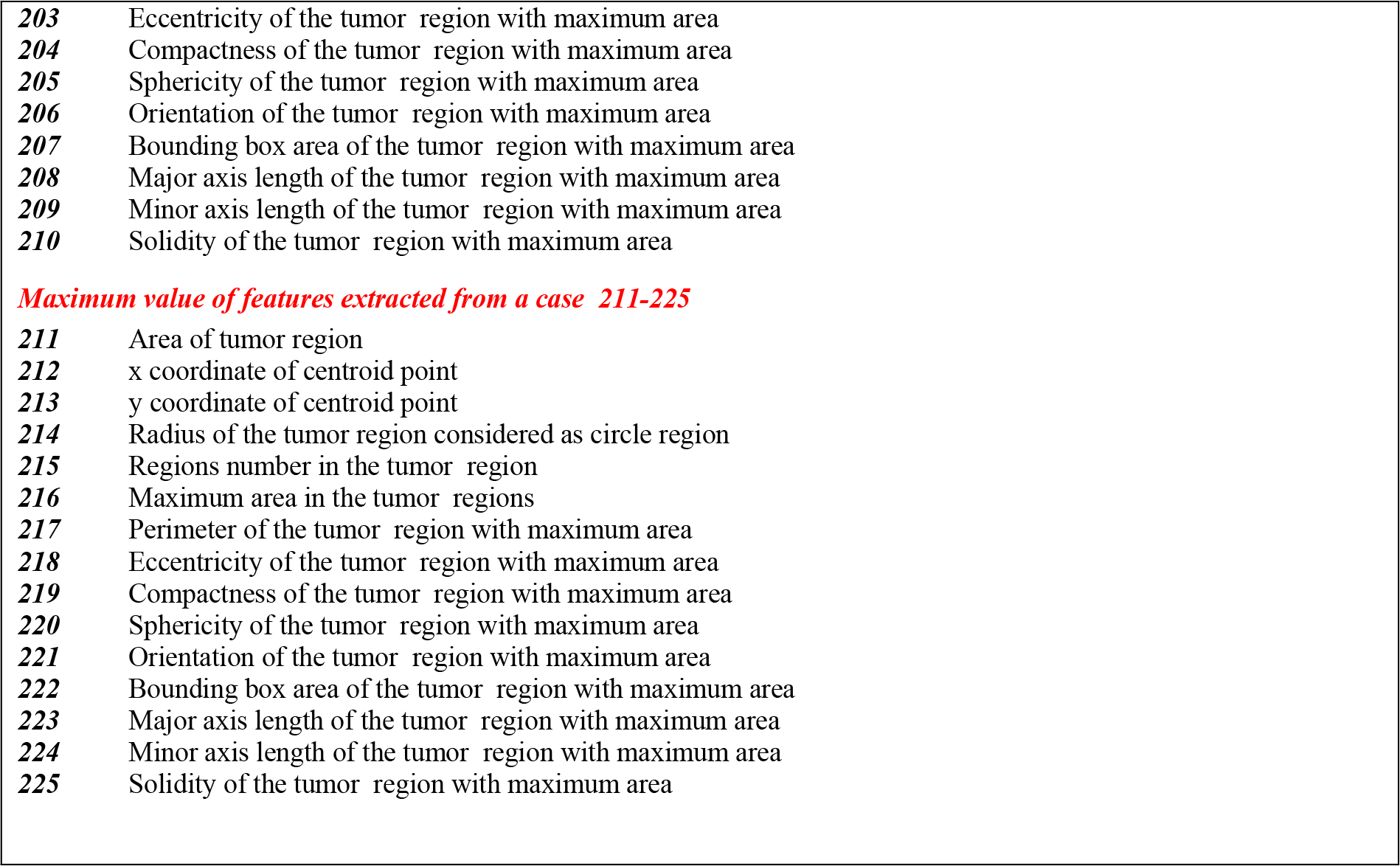

**Table S2.**
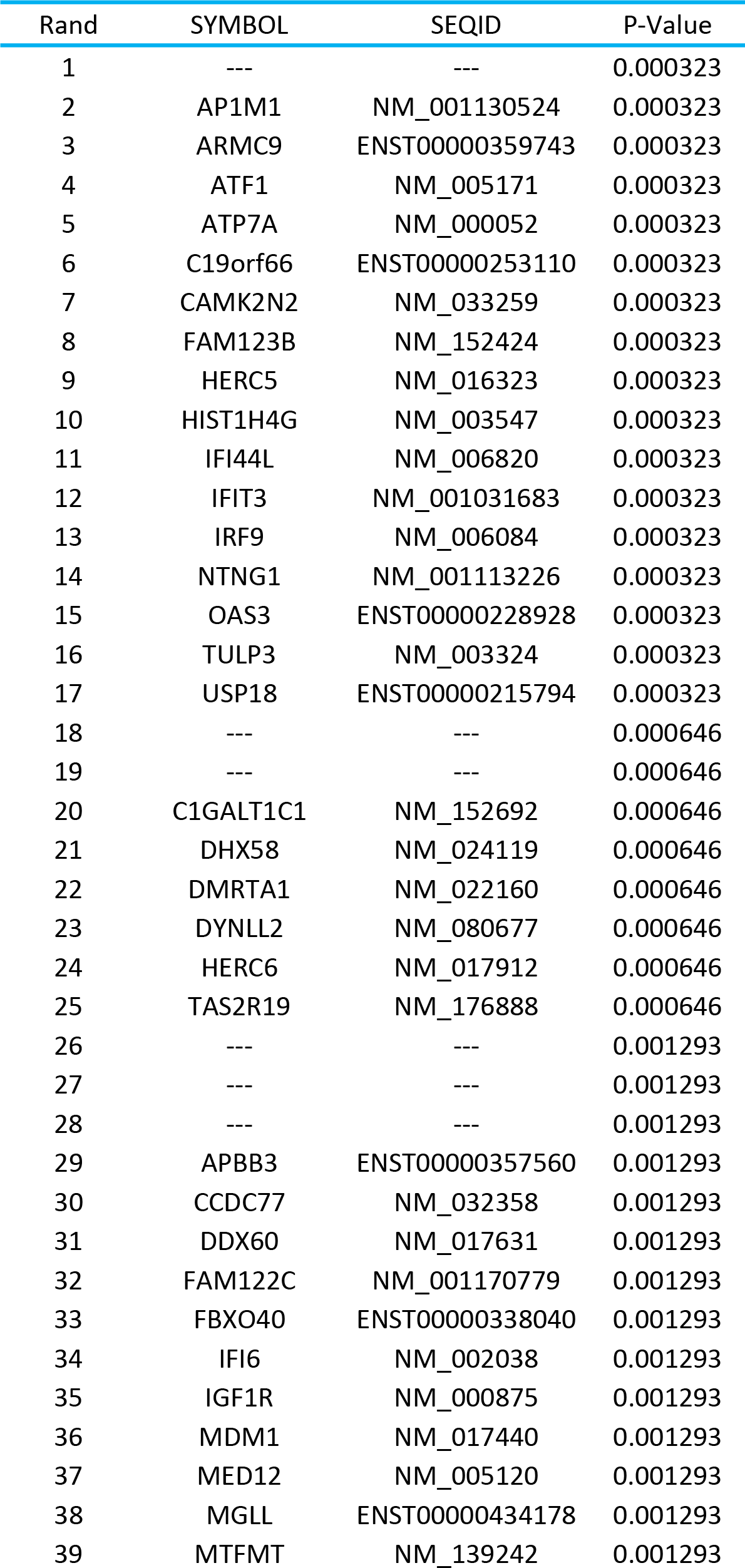
Top 119 significant genes screened by the Wilcoxon rank sum test with P-value of 0.005.

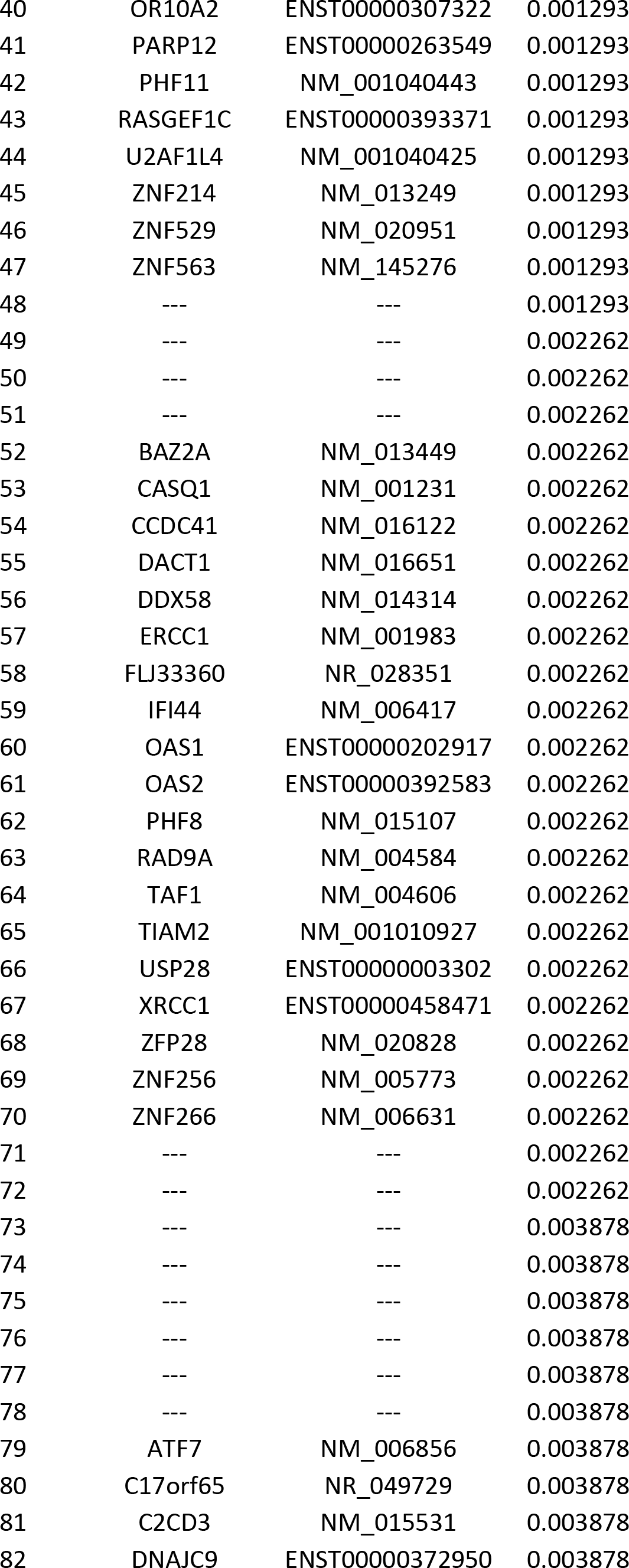

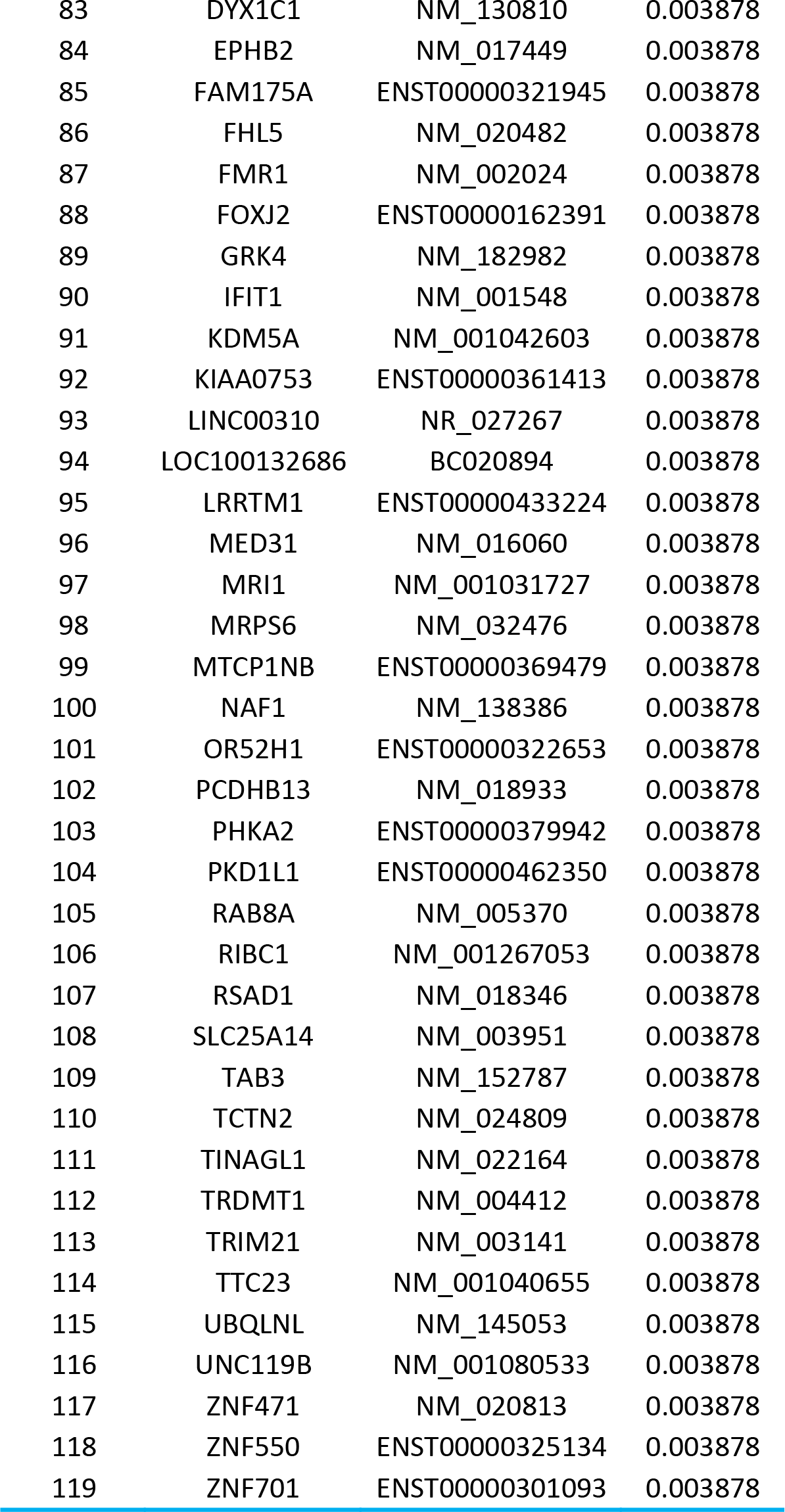

|  |  |  |  |
| --- | --- | --- | --- |
| 40 | OR10A2 | ENST00000307322 | 0.001293 |
| 41 | PARP12 | ENST00000263549 | 0.001293 |
| 42 | PHF11 | NM_001040443 | 0.001293 |
| 43 | RASGEF1C | ENST00000393371 | 0.001293 |
| 44 | U2AF1L4 | NM_001040425 | 0.001293 |
| 45 | ZNF214 | NM_013249 | 0.001293 |
| 46 | ZNF529 | NM_020951 | 0.001293 |
| 47 | ZNF563 | NM_145276 | 0.001293 |
| 48 | --- | --- | 0.001293 |
| 49 | --- | --- | 0.002262 |
| 50 | --- | --- | 0.002262 |
| 51 | --- | --- | 0.002262 |
| 52 | BAZ2A | NM_013449 | 0.002262 |
| 53 | CASQ1 | NM_001231 | 0.002262 |
| 54 | CCDC41 | NM_016122 | 0.002262 |
| 55 | DACT1 | NM_016651 | 0.002262 |
| 56 | DDX58 | NM_014314 | 0.002262 |
| 57 | ERCC1 | NM_001983 | 0.002262 |
| 58 | FLJ33360 | NR_028351 | 0.002262 |
| 59 | IFI44 | NM_006417 | 0.002262 |
| 60 | OAS1 | ENST00000202917 | 0.002262 |
| 61 | OAS2 | ENST00000392583 | 0.002262 |
| 62 | PHF8 | NM_015107 | 0.002262 |
| 63 | RAD9A | NM_004584 | 0.002262 |
| 64 | TAF1 | NM_004606 | 0.002262 |
| 65 | TIAM2 | NM_001010927 | 0.002262 |
| 66 | USP28 | ENST00000003302 | 0.002262 |
| 67 | XRCC1 | ENST00000458471 | 0.002262 |
| 68 | ZFP28 | NM_020828 | 0.002262 |
| 69 | ZNF256 | NM_005773 | 0.002262 |
| 70 | ZNF266 | NM_006631 | 0.002262 |
| 71 | --- | --- | 0.002262 |
| 72 | --- | --- | 0.002262 |
| 73 | --- | --- | 0.003878 |
| 74 | --- | --- | 0.003878 |
| 75 | --- | --- | 0.003878 |
| 76 | --- | --- | 0.003878 |
| 77 | --- | --- | 0.003878 |
| 78 | --- | --- | 0.003878 |
| 79 | ATF7 | NM_006856 | 0.003878 |
| 80 | C17orf65 | NR_049729 | 0.003878 |
| 81 | C2CD3 | NM_015531 | 0.003878 |
| 82 | DNAJC9 | ENST00000372950 | 0.003878 |
| 83 | DYX1C1 | NM_130810 | 0.003878 |
| 84 | EPHB2 | NM_017449 | 0.003878 |
| 85 | FAM175A | ENST00000321945 | 0.003878 |
| 86 | FHL5 | NM_020482 | 0.003878 |
| 87 | FMR1 | NM_002024 | 0.003878 |
| 88 | FOXJ2 | ENST00000162391 | 0.003878 |
| 89 | GRK4 | NM_182982 | 0.003878 |
| 90 | IFIT1 | NM_001548 | 0.003878 |
| 91 | KDM5A | NM_001042603 | 0.003878 |
| 92 | KIAA0753 | ENST00000361413 | 0.003878 |
| 93 | LINC00310 | NR_027267 | 0.003878 |
| 94 | LOC100132686 | BC020894 | 0.003878 |
| 95 | LRRTM1 | ENST00000433224 | 0.003878 |
| 96 | MED31 | NM_016060 | 0.003878 |
| 97 | MRI1 | NM_001031727 | 0.003878 |
| 98 | MRPS6 | NM_032476 | 0.003878 |
| 99 | MTCP1NB | ENST00000369479 | 0.003878 |
| 100 | NAF1 | NM_138386 | 0.003878 |
| 101 | OR52H1 | ENST00000322653 | 0.003878 |
| 102 | PCDHB13 | NM_018933 | 0.003878 |
| 103 | PHKA2 | ENST00000379942 | 0.003878 |
| 104 | PKD1L1 | ENST00000462350 | 0.003878 |
| 105 | RAB8A | NM_005370 | 0.003878 |
| 106 | RIBC1 | NM_001267053 | 0.003878 |
| 107 | RSAD1 | NM_018346 | 0.003878 |
| 108 | SLC25A14 | NM_003951 | 0.003878 |
| 109 | TAB3 | NM_152787 | 0.003878 |
| 110 | TCTN2 | NM_024809 | 0.003878 |
| 111 | TINAGL1 | NM_022164 | 0.003878 |
| 112 | TRDMT1 | NM_004412 | 0.003878 |
| 113 | TRIM21 | NM_003141 | 0.003878 |
| 114 | TTC23 | NM_001040655 | 0.003878 |
| 115 | UBQLNL | NM_145053 | 0.003878 |
| 116 | UNC119B | NM_001080533 | 0.003878 |
| 117 | ZNF471 | NM_020813 | 0.003878 |
| 118 | ZNF550 | ENST00000325134 | 0.003878 |
| 119 | ZNF701 | ENST00000301093 | 0.003878 |

**Table S3.**
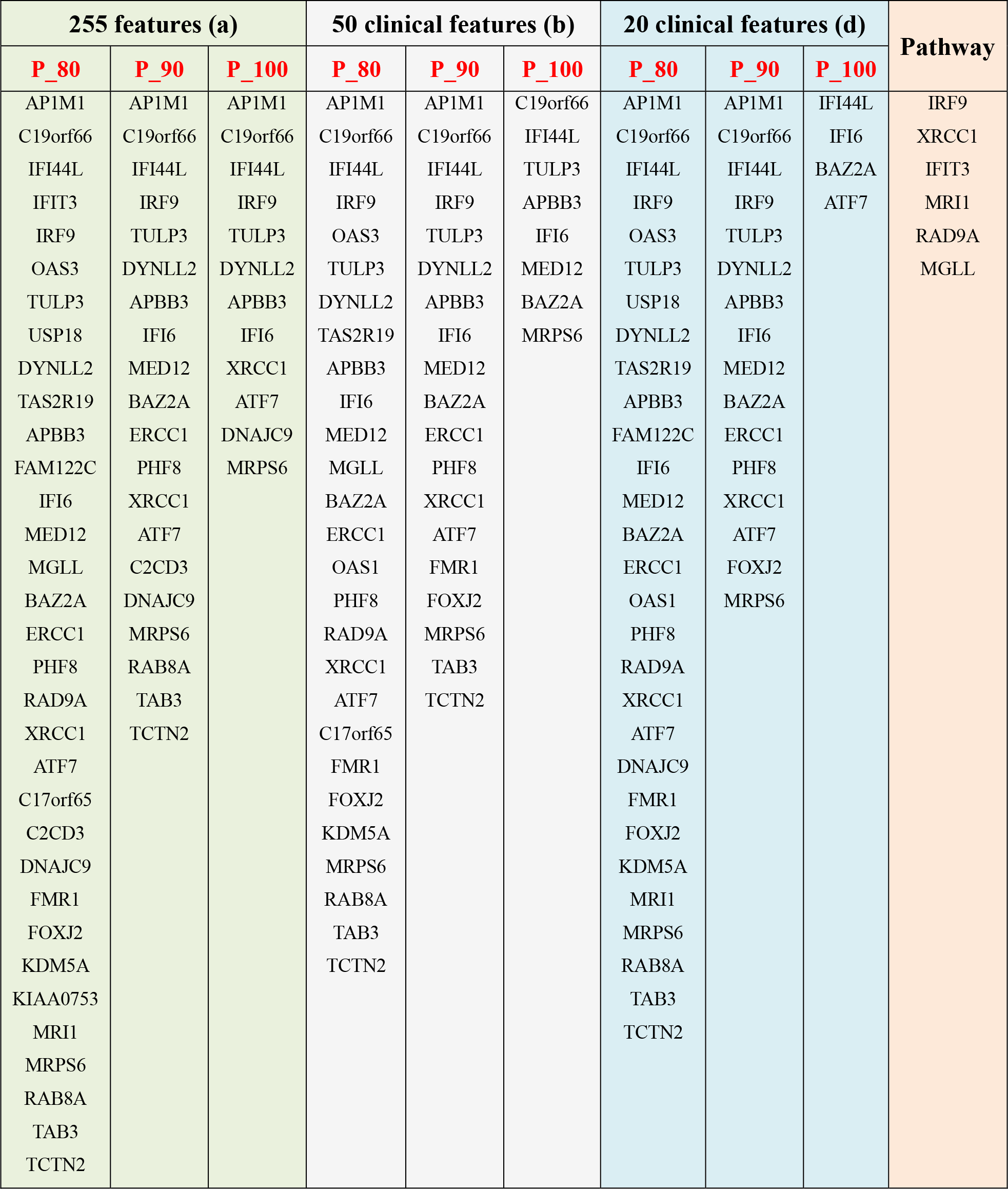
The candidate genes selected from three representative feature sets of 255 features, 50 clinical features and 20 clinical features (corresponding to the Fig. S5 (a), (b) and (d), respectively). *P_100, P_90 and P_80* mean 100%, 90% and 80% of coverage *P*, respectively. The genes from top ranked canonical pathways were in the right column.

| 255 features (a) |  |  | 50 clinical features (b) |  |  | 20 clinical features (d) |  |  | Pathway |
| --- | --- | --- | --- | --- | --- | --- | --- | --- | --- |
| $P_{80}$ | $P_{90}$ | $P_{100}$ | $P_{80}$ | $P_{90}$ | $P_{100}$ | $P_{80}$ | $P_{90}$ | $P_{100}$ | |
| AP1M1 | AP1M1 | AP1M1 | AP1M1 | AP1M1 | C19orf66 | AP1M1 | AP1M1 | IFI44L | IRF9 |
| C19orf66 | C19orf66 | C19orf66 | C19orf66 | C19orf66 | IFI44L | C19orf66 | C19orf66 | IFI6 | XRCC1 |
| IFI44L | IFI44L | IFI44L | IFI44L | IFI44L | TULP3 | IFI44L | IFI44L | BAZ2A | IFIT3 |
| IFIT3 | IRF9 | IRF9 | IRF9 | IRF9 | APBB3 | IRF9 | IRF9 | ATF7 | MRI1 |
| IRF9 | TULP3 | TULP3 | OAS3 | TULP3 | IFI6 | OAS3 | TULP3 |  | RAD9A |
| OAS3 | DYNLL2 | DYNLL2 | TULP3 | DYNLL2 | MED12 | TULP3 | DYNLL2 |  | MGLL |
| TULP3 | APBB3 | APBB3 | DYNLL2 | APBB3 | BAZ2A | USP18 | APBB3 |  |  |
| USP18 | IFI6 | IFI6 | TAS2R19 | IFI6 | MRPS6 | DYNLL2 | IFI6 |  |  |
| DYNLL2 | MED12 | XRCC1 | APBB3 | MED12 |  | TAS2R19 | MED12 |  |  |
| TAS2R19 | BAZ2A | ATF7 | IFI6 | BAZ2A |  | APBB3 | BAZ2A |  |  |
| APBB3 | ERCC1 | DNAJC9 | MED12 | ERCC1 |  | FAM122C | ERCC1 |  |  |
| FAM122C | PHF8 | MRPS6 | MGLL | PHF8 |  | IFI6 | PHF8 |  |  |
| IFI6 | XRCC1 |  | BAZ2A | XRCC1 |  | MED12 | XRCC1 |  |  |
| MED12 | ATF7 |  | ERCC1 | ATF7 |  | BAZ2A | ATF7 |  |  |
| MGLL | C2CD3 |  | OAS1 | FMR1 |  | ERCC1 | FOXJ2 |  |  |
| BAZ2A | DNAJC9 |  | PHF8 | FOXJ2 |  | OAS1 | MRPS6 |  |  |
| ERCC1 | MRPS6 |  | RAD9A | MRPS6 |  | PHF8 |  |  |  |
| PHF8 | RAB8A |  | XRCC1 | TAB3 |  | RAD9A |  |  |  |
| RAD9A | TAB3 |  | ATF7 | TCTN2 |  | XRCC1 |  |  |  |
| XRCC1 | TCTN2 |  | C17orf65 |  |  | ATF7 |  |  |  |
| ATF7 |  |  | FMR1 |  |  | DNAJC9 |  |  |  |
| C17orf65 |  |  | FOXJ2 |  |  | FMR1 |  |  |  |
| C2CD3 |  |  | KDM5A |  |  | FOXJ2 |  |  |  |
| DNAJC9 |  |  | MRPS6 |  |  | KDM5A |  |  |  |
| FMR1 |  |  | RAB8A |  |  | MRI1 |  |  |  |
| FOXJ2 |  |  | TAB3 |  |  | MRPS6 |  |  |  |
| KDM5A |  |  | TCTN2 |  |  | RAB8A |  |  |  |
| KIAA0753 |  |  |  |  |  | TAB3 |  |  |  |
| MRI1 |  |  |  |  |  | TCTN2 |  |  |  |
| MRPS6 |  |  |  |  |  |  |  |  |  |
| RAB8A |  |  |  |  |  |  |  |  |  |
| TAB3 |  |  |  |  |  |  |  |  |  |
| TCTN2 |  |  |  |  |  |  |  |  |  |

**Table S4.** Predication of MGMT methylation status in previous studies

|  |  | Methylated MGMT |  | Unmethylated MGMT |  | Sensitivity <sup>(1)</sup> | Specificity |
| --- | --- | --- | --- | --- | --- | --- | --- |
| Brandes, et al<sup>1</sup> | PsP | 21 | 91.3% | 11 | 40.7% | 65.6% | 88.9% |
|  | TTP | 2 | 8.7% | 16 | 59.3% |  |  |
| Motegi, et al<sup>2</sup> | PsP | 2 | 40% | 1 | 16.7% | 66.7% | 62.5% |
|  | TTP | 3 | 60% | 5 | 83.3% |  |  |
| Kong, et al<sup>3</sup> | PsP | 15 | 37.5% | 8 | 24.2% | 65.1% | 50% |
|  | TTP | 25 | 62.5% | 25 | 75.8% |  |  |
| Mieghem, et al<sup>4</sup> (2) | PsP | 9 | 37.5% | 4 | 12.9% | 69.2% | 64.3% |
|  | TTP | 15 | 62.5% | 27 | 87.1% |  |  |
| Park, et al<sup>5</sup> (3) | PsP | 4 | 80% | 7 | 35% | 36.4% | 92.9% |
|  | TTP | 1 | 20% | 13 | 65% |  |  |

(1): Sensitivity and specificity were defined for detecting the PsP based on methylated MGMT; (2): MGMT assessed using liberal criteria; (3) MGMT promoter methylation status determined by methylation-specific polymerase chain reaction.

**Table S5.**
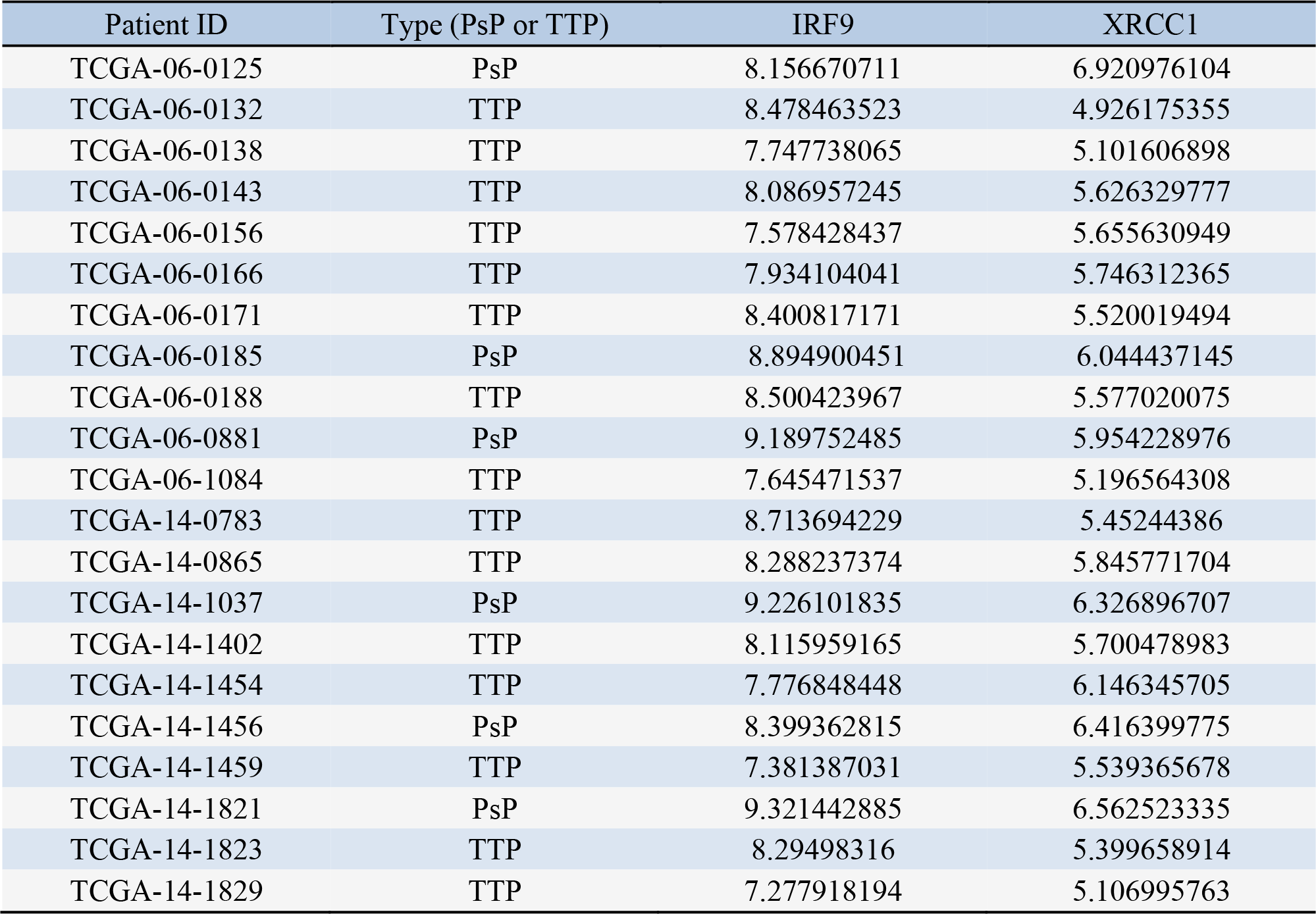
Gene expressions of IRF9 and XRCC1 in the samples from TCGA. The sample type, i.e., pseudoprogression (PsP) or true tumor progression (TTP), is determined by the longitudinal MRI from TCIA.

## Comparison with classic scheme (i.e. directly using the sample labels for marker identification)

We have compared our approach with the classic scheme (i.e. directly using the sample labels for marker identification), as shown in supplementary Fig. S2 and S3. Actually, our approach also adopted the phenotype labels. Specifically, in the first step, we compared the gene expression levels in the PsP and TTP groups to obtain the differentially expressed genes. We initially identified 119 genes using the Wilcoxon rank sum test with p<0.005. After that, we selected 33 candidate genes using associations study between imaging features and gene expression profiles with two advantages: 1) the imaging features were extracted from tumors along the longitudinal MRI and provided diagnostic information of PsP and TTP. Thus, the 33 candidate genes were confirmed to be associated with the development of PsP and TTP. 2) The association study can narrow down candidate genes from 119 to 33, which improves the efficiency of the biological relevance analysis. In summary, the association study based on the differentially expressed genes facilitates the biological analysis in efficiently identifying most relevant signaling pathways.

To illustrate the advantage of our approach, we conducted the comparison experiments (Fig. S2 and S3). First, we only used the phenotype labels to obtain 119 differentially expressed genes using the Wilcoxon rank sum test with p<0.005. We then performed the biological analysis for the 119 genes without the association study. Top-ranked Canonical Pathways and their genes by IPA were shown in Fig. S3. Obviously, the XRCC1 and its pathways, such as BER pathway, DNA Double-strand Break Repair by Non-Homologous End Joining, and DNA damage-induced 14-3-3σsignaling, in the top-ranked lists of our study (Association study, Fig. S2) were absent from the significant lists of the classic scheme (Fig. S3 (a)). The p-value of these XRCC1 pathways from the classic scheme was slightly greater than 0.05 (Fig. S3 (b)). Other pathways in the top-ranked list, such as the Role of RIG1-like Receptors in Antiviral innate immunity, the Role of BRCA1 in DAN damage response, and the role of pattern recognition receptors in recognition of bacteria and viruses, were not directly related to cancer development. This result indicates that differentially expressed genes without relation to the PsP and TTP may confound the biological analysis. Therefore, the association study can not only confirm the candidate genes with relation to the development of PsP and TTP but also narrow down the candidate genes, thereby ensuring the real significant genes stand out in the biological analysis.

## Effect of morphological features with different size

To illustrate the effect of features, we conducted comparisons of six feature sets with different sizes, as shown in Fig. S5. The six feature sets were 255 morphological features, 50 clinical morphological features, 50 features randomly selected from our 255 features, and 20 features randomly selected from the 50 clinical features for three times, corresponding to Figure S5 (a)-(f), respectively. Specifically, we selected 50 clinical morphological features from 255 features, corresponding to the VASARI, which offers a set of well-defined terms that describe the GBM tumors^1^. The 50 clinical morphological features contain the major/minor axis length of the enhanced region, the thickness of enhancing margin, tumor volume, proportion enhancing and proportion necrosis, etc. In each experiment of the individual feature set, there were 27 parameter combinations for our model. We defined the coverage rate *P* as the occurrence frequency of genes on the lists of weight ranked top 50 in 27 parameter combinations. As a result, we can obtain three different sets of candidate genes with 80%, 90% and 100% of coverage *P* for the individual feature set. Results from three representative feature sets of 255 features, 50 clinical features, and 20 clinical features (corresponding to the Fig. S5 (a), (b) and (d), respectively) were shown in Table S3. Then, the candidate genes identified by coverage *P=0.8* were used for pathway analysis using IPA. Not surprisingly, the genes from the top-ranked canonical pathways from six feature sets, called pathway genes, were same, as shown in the right column of Table S3. Fig. S5 shows the gene overlap among candidate genes from different coverage *P* and pathway genes. Obviously, when the feature number is 20 or 50, there is no overlap between candidate genes from *P=100%* and pathway genes. Conversely, our model with 255 features yielded the biomarkers, i.e., IRF9 and XRCC1, in all of the three coverage rates. These results indicate that the performance of our model with 255 features is more robust than that with 50 or 20 features.

## Reference

1. Bleeker, F.E., Molenaar, R.J. & Leenstra, S. Recent advances in the molecular understanding of glioblastoma. J Neurooncol 108, 11–27 (2012).

2. Brandsma, D., Stalpers, L., Taal, W., Sminia, P. & van den Bent, M.J. Clinical features, mechanisms, and management of pseudoprogression in malignant gliomas. The lancet oncology 9, 453–461 (2008).

3. Kruser, T.J., Mehta, M.P. & Robins, H.I. Pseudoprogression after glioma therapy: a comprehensive review. Expert Rev Neurother 13, 389–403 (2013).

4. Gerstner, E.R., McNamara, M.B., Norden, A.D., Lafrankie, D. & Wen, P.Y. Effect of adding temozolomide to radiation therapy on the incidence of pseudo-progression. J Neurooncol 94, 97–101 (2009).

5. Taal, W. et al. Incidence of early pseudo-progression in a cohort of malignant glioma patients treated with chemoirradiation with temozolomide. Cancer 113, 405–10 (2008).

6. Bisdas, S. et al. Cerebral Blood Volume Measurements by Perfusion-Weighted MR Imaging in Gliomas: Ready for Prime Time in Predicting Short-Term Outcome and Recurrent Disease? American Journal of Neuroradiology 30, 681–688 (2009).

7. Law, M. et al. Gliomas: Predicting time to progression or survival with cerebral blood volume measurements at dynamic susceptibility-weighted contrast-enhanced perfusion MR imaging. Radiology 247, 490–498 (2008).

8. Jain, R. et al. Genomic Mapping and Survival Prediction in Glioblastoma: Molecular Subclassification Strengthened by Hemodynamic Imaging Biomarkers. Radiology 267, 212–220 (2013).

9. Hirai, T. et al. Prognostic value of perfusion MR imaging of high-grade astrocytomas: Long-term follow-up study. American Journal of Neuroradiology 29, 1505–1510 (2008).

10. Park, J.K. et al. Scale to Predict Survival After Surgery for Recurrent Glioblastoma Multiforme. Journal of Clinical Oncology 28, 3838–3843 (2010).

11. Gutman, D.A. et al. MR Imaging Predictors of Molecular Profile and Survival: Multi-institutional Study of the TCGA Glioblastoma Data Set. Radiology 267, 560–569 (2013).

12. Brandes, A.A. et al. MGMT promoter methylation status can predict the incidence and outcome of pseudoprogression after concomitant radiochemotherapy in newly diagnosed glioblastoma patients. J Clin Oncol 26, 2192–7 (2008).

13. Pouleau, H.B. et al. High levels of cellular proliferation predict pseudoprogression in glioblastoma patients. Int J Oncol 40, 923–8 (2012).

14. Motegi, H. et al. IDH1 mutation as a potential novel biomarker for distinguishing pseudoprogression from true progression in patients with glioblastoma treated with temozolomide and radiotherapy. Brain Tumor Pathol 30, 67–72 (2013).

15. Kang, H.C. et al. Pseudoprogression in patients with malignant gliomas treated with concurrent temozolomide and radiotherapy: potential role of p53. J Neurooncol 102, 157–62 (2011).

16. Melguizo-Gavilanes, I., Bruner, J.M., Guha-Thakurta, N., Hess, K.R. & Puduvalli, V.K. Characterization of pseudoprogression in patients with glioblastoma: is histology the gold standard? J Neurooncol 123, 141–50 (2015).

17. Hayes, J. et al. Prediction of clinical outcome in glioblastoma using a biologically relevant nine-microRNA signature. Mol Oncol 9, 704–14 (2015).

18. da Cruz, L.C.H., Rodriguez, I., Domingues, R.C., Gasparetto, E.L. & Sorensen, A.G. Pseudoprogression and Pseudoresponse: Imaging Challenges in the Assessment of Posttreatment Glioma. American Journal of Neuroradiology 32, 1978–1985 (2011).

19. Van Mieghem, E. et al. Defining pseudoprogression in glioblastoma multiforme. Eur J Neurol 20, 1335–41 (2013).

20. Pinho, M.C. et al. Low Incidence of Pseudoprogression by Imaging in Newly Diagnosed Glioblastoma Patients Treated With Cediranib in Combination With Chemoradiation. Oncologist 19, 75–81 (2014).

21. Kong, D.S. et al. Diagnostic dilemma of pseudoprogression in the treatment of newly diagnosed glioblastomas: the role of assessing relative cerebral blood flow volume and oxygen-6-methylguanine-DNA methyltransferase promoter methylation status. AJNR Am J Neuroradiol 32, 382–7 (2011).

22. Park, C.K. et al. Usefulness of MS-MLPA for detection of MGMT promoter methylation in the evaluation of pseudoprogression in glioblastoma patients. Neuro Oncol 13, 195–202 (2011).

23. Topkan, E., Topuk, S., Oymak, E., Parlak, C. & Pehlivan, B. Pseudoprogression in patients with glioblastoma multiforme after concurrent radiotherapy and temozolomide. Am J Clin Oncol 35, 284–9 (2012).

24. Nasseri, M. et al. Evaluation of pseudoprogression in patients with glioblastoma multiforme using dynamic magnetic resonance imaging with ferumoxytol calls RANO criteria into question. Neuro Oncol 16, 1146–54 (2014).

25. Kuo, M.D. & Jamshidi, N. Behind the numbers: Decoding molecular phenotypes with radiogenomics--guiding principles and technical considerations. Radiology 270, 320–5 (2014).

26. Beck, A.H. et al. Systematic Analysis of Breast Cancer Morphology Uncovers Stromal Features Associated with Survival. Science Translational Medicine 3(2011).

27. Aziz, N.A.A. et al. A 19-Gene expression signature as a predictor of survival in colorectal cancer. Bmc Medical Genomics 9(2016).

28. van de Vijver, M.J. et al. A gene-expression signature as a predictor of survival in breast cancer. New England Journal of Medicine 347, 1999–2009 (2002).

29. Parvez, K., Parvez, A. & Zadeh, G. The diagnosis and treatment of pseudoprogression, radiation necrosis and brain tumor recurrence. Int J Mol Sci 15, 11832–46 (2014).

30. Verma, N., Cowperthwaite, M.C., Burnett, M.G. & Markey, M.K. Differentiating tumor recurrence from treatment necrosis: a review of neuro-oncologic imaging strategies. Neuro Oncol 15, 515–34 (2013).

31. Qian, X. et al. Stratification of Pseudoprogression and True Progression of GBM based on longitudinal DTI without Segmentation. Medical Physics (2016).

32. Qian, X. et al. Objective classification system for sagittal craniosynostosis based on suture segmentation. Med Phys 42, 5545 (2015).

33. Tan, H., Bao, J. & Zhou, X. A novel missense-mutation-related feature extraction scheme for ‘driver’ mutation identification. Bioinformatics 28, 2948–55 (2012).

34. Chang, C.-C., and Chih-Jen Lin. LIBSVM: a library for support vector machines. ACM Transactions on Intelligent Systems and Technology 2, 21–27 (2011).

35. Qian, X. et al. Identification of biomarkers for pseudo and true progression of GBM based on radiogenomics study. Oncotarget (2016).

36. Riemenschneider, M.J., Jeuken, J.W.M., Wesseling, P. & Reifenberger, G. Molecular diagnostics of gliomas: state of the art. Acta Neuropathologica 120, 567–584 (2010).

37. Touat, M. et al. Emerging circulating biomarkers in glioblastoma: promises and challenges. Expert Review of Molecular Diagnostics 15, 1311–1323 (2015).

38. Takaoka, A. et al. Integration of interferon-alpha/beta signalling to p53 responses in tumour suppression and antiviral defence. Nature 424, 516–523 (2003).

39. Wang, P.X. et al. Interferon regulatory factor 9 is a key mediator of hepatic ischemia/reperfusion injury. Journal of Hepatology 62, 111–120 (2015).

40. Takaoka, A., Tamura, T. & Taniguchi, T. Interferon regulatory factor family of transcription factors and regulation of oncogenesis. Cancer Science 99, 467–478 (2008).

41. Lau, J.F., Parisien, J.P. & Horvath, C.M. Interferon regulatory factor subcellular localization is determined by a bipartite nuclear localization signal in the DNA-binding domain and interaction with cytoplasmic retention factors. Proceedings of the National Academy of Sciences of the United States of America 97, 7278–7283 (2000).

42. Tsuno, T. et al. IRF9 is a Key Factor for Eliciting the Antiproliferative Activity of IFN-alpha. Journal of Immunotherapy 32, 803–816 (2009).

43. Yanai, H., Negishi, H. & Taniguchi, T. The IRF family of transcription factors Inception, impact and implications in oncogenesis. Oncoimmunology 1, 1376–1386 (2012).

44. Luker, K.E., Pica, C.M., Schreiber, R.D. & Piwnica-Worms, D. Overexpression of IRF9 confers resistance to antimicrotubule agents in breast cancer cells. Cancer Research 61, 6540–6547 (2001).

45. Gong, X. et al. A brain-wide association study of DISC1 genetic variants reveals a relationship with the structure and functional connectivity of the precuneus in schizophrenia. Hum Brain Mapp 35, 5414–30 (2014).

46. Aerts, H.J. et al. Decoding tumour phenotype by noninvasive imaging using a quantitative radiomics approach. Nat Commun 5, 4006 (2014).

47. Yuan, Y. et al. Quantitative image analysis of cellular heterogeneity in breast tumors complements genomic profiling. Sci Transl Med 4, 157ra143 (2012).

## Reference

1. Thomas Jefferson University hospital, “VASARI MRI visual feature guide” (2010)

## Reference

1 A.A. Brandes, E. Franceschi, A. Tosoni, V. Blatt, A. Pession, G. Tallini, R. Bertorelle, S. Bartolini, F. Calbucci, A. Andreoli, G. Frezza, M. Leonardi, F. Spagnolli, M. Ermani, “MGMT promoter methylation status can predict the incidence and outcome of pseudoprogression after concomitant radiochemotherapy in newly diagnosed glioblastoma patients,” Journal of Clinical Oncology 26, 2192–2197 (2008).

2 H. Motegi, Y. Kamoshima, S. Terasaka, H. Kobayashi, S. Yamaguchi, M. Tanino, J. Murata, K. Houkin, “IDH1 mutation as a potential novel biomarker for distinguishing pseudoprogression from true progression in patients with glioblastoma treated with temozolomide and radiotherapy,” Brain Tumor Pathol 30, 67–72 (2013).

3 D.S. Kong, S.T. Kim, E.H. Kim, D.H. Lim, W.S. Kim, Y.L. Suh, J.I. Lee, K. Park, J.H. Kim, D.H. Nam, “Diagnostic dilemma of pseudoprogression in the treatment of newly diagnosed glioblastomas: the role of assessing relative cerebral blood flow volume and oxygen-6-methylguanine-DNA methyltransferase promoter methylation status,” AJNR Am J Neuroradiol 32, 382–387 (2011).

4 E. Van Mieghem, A. Wozniak, Y. Geussens, J. Menten, S. De Vleeschouwer, F. Van Calenbergh, R. Sciot, S. Van Gool, O.E. Bechter, P. Demaerel, G. Wilms, P.M. Clement, “Defining pseudoprogression in glioblastoma multiforme,” Eur J Neurol 20, 1335–1341 (2013).

5 C.K. Park, J. Kim, S.Y. Yim, A.R. Lee, J.H. Han, C.Y. Kim, S.H. Park, T.M. Kim, S.H. Lee, S.H. Choi, S.K. Kim, D.G. Kim, H.W. Jung, “Usefulness of MS-MLPA for detection of MGMT promoter methylation in the evaluation of pseudoprogression in glioblastoma patients,” Neuro Oncol 13, 195–202 (2011).

